# Blood and cerebellar abundance of *ATXN3* splice variants in spinocerebellar ataxia type 3/Machado-Joseph disease

**DOI:** 10.1101/2023.04.22.537936

**Authors:** Mafalda Raposo, Jeannette Hübener-Schmid, Rebecca Tagett, Ana F. Ferreira, Ana Rosa Vieira Melo, João Vasconcelos, Paula Pires, Teresa Kay, Hector Garcia-Moreno, Paola Giunti, Magda M. Santana, Luis Pereira de Almeida, Jon Infante, Bart P. van de Warrenburg, Jeroen J. de Vries, Jennifer Faber, Thomas Klockgether, Nicolas Casadei, Jakob Admard, Ludger Schöls, Olaf Riess, European Spinocerebellar ataxia type 3/Machado-Joseph disease Initiative (ESMI) study group, Maria do Carmo Costa, Manuela Lima

## Abstract

**Background:** Spinocerebellar ataxia type 3 (SCA3)/Machado-Joseph disease (MJD) is an autosomal dominant polyglutamine disease. SCA3/MJD causative gene, *ATXN3*, is known to undergo alternative splicing (AS) and 54 transcripts are currently annotated. Differences in the toxicity of ataxin-3 protein isoforms, harbouring on its C-terminus two or three ubiquitin interacting motifs (UIMs), were previously uncovered, raising the hypothesis that specific *ATXN3* splice variants play key roles in promoting the selective toxicity displayed in SCA3/MJD.

**Methods:** Using RNA-seq datasets we identified and determined the abundance of annotated *ATXN3* transcripts in blood (n=60) and cerebellum (n=12) of SCA3/MJD subjects and controls.

**Results:** Globally, the number and the abundance of individual *ATXN3* transcripts were higher in the cerebellum than in the blood. While the most abundant transcript in the cerebellum was a protein with a coding sequence not defined of unknown function (*ATXN3*-208), the transcript with the highest abundance in blood was the reference transcript (*ATXN3*-251) which translates into an ataxin-3 isoform harboring three UIMs. Noteworthy, the abundance of *ATXN3*-251 and *ATXN3*-214, two out of the four transcripts that encode full-length ataxin-3 protein isoforms but differ in the C-terminus were strongly related with tissue expression specificity: *ATXN3-251* (3UIM) was expressed in blood 50-fold more than in cerebellum, whereas *ATXN3*-214 (2UIM) was expressed in the cerebellum 20-fold more than in blood.

**Conclusions:** These findings provide new insights into the elucidation of *ATXN3* AS in different tissues, contributing for a better understanding of SCA3/MJD pathogenesis and providing information for the development of future effective *ATXN3* mRNA-lowering therapies.

## INTRODUCTION

Alternative splicing (AS) is a fundamental step in the regulation of eukaryotic gene expression through which introns are excised and exons are ligated together to form the messenger RNA (mRNA) (Vuong *et al*, 2016). Multiple evidence indicates that genetic variation in *cis*-acting splicing sequence elements or in *trans*-acting splicing factors plays a relevant role in patients with neurodegenerative conditions (reviewed in (Daguenet *et al*, 2015; Nik & Bowman, 2019)). Importantly, *post-mortem* brains of Huntington disease (HD) patients and of mouse models show aberrant splicing of the HD causative gene (huntingtin, *HTT*) itself, leading to the production of high levels of a splice variant only containing exon 1 that encodes a highly aggregation-prone protein and thus contributes to disease pathogenesis (Mangiarini *et al*, 1996; Sathasivam *et al*, 2013; Barbaro *et al*, 2015; Neueder *et al*, 2017). Currently, several pathologies triggered by splicing defects, including neurodegenerative diseases, can be treated using different types of RNA molecules, such as antisense oligonucleotide (ASO) or small interference RNA (siRNA), either by reducing the levels of the defective protein or by producing the corrected protein (reviewed in (Dhuri *et al*, 2020; Zhu *et al*, 2022)). In fact, ASO-mediated therapies are currently available for some common and rare diseases, including neurological diseases such as spinal muscular atrophy and Duchenne muscular dystrophy (reviewed in (Zhu *et al*, 2022)).

SCA3/MJD is an autosomal dominant neurodegenerative polyglutamine (polyQ) disorder and the most common dominant ataxia worldwide (Sequeiros *et al*, 2012). SCA3/MJD is caused by an abnormal polyQ-encoding CAG repeat motif, consensually containing more than 60 trinucleotides within exon 10 of the *ATXN3* gene (Takiyama *et al*, 1993; Kawaguchi *et al*, 1994; Maciel *et al*, 2001). The *ATXN3* gene (ENSG00000066427) originates 54 transcripts (Martin *et al*, 2023) in the GRCh38.p13 human genome assembly including the canonical transcript (ENST00000644486/*ATXN3*-251) containing 11 exons and spanning a genomic region about 48 kb (Ichikawa *et al*, 2001), encoding the ataxin-3 canonical protein isoform (P54252-2, UniProt).

Ataxin-3 is a ubiquitously expressed deubiquitinating enzyme (*e.g.*, (Burnett *et al*, 2003)) harbouring on its N-terminus the globular Josephin domain (JD) that includes the catalytic residues, and on its C-terminus a protein segment of undetermined structure containing the polyQ tract and two or three ubiquitin interacting motifs (UIM) (Goto *et al*, 1997; Masino *et al*, 2003). Aggregation of the ataxin-3 protein is a hallmark of SCA3/MJD (reviewed in (Costa & Paulson, 2012)) and occurs not only in brain regions with neuronal loss, such as the cerebellar dentate nucleus and the motor and pontine nuclei of the brainstem, but also in brain areas spared of neuronal destruction, like the cerebellar cortex (Koeppen, 2018).

Despite nearly 30 years of research following the identification of *ATXN3* as the causative gene of SCA3/MJD (Takiyama *et al*, 1993; Kawaguchi *et al*, 1994), the characterization of the transcriptional species of *ATXN3* remains very incomplete. While *ATXN3* is known to undergo AS (Kawaguchi *et al*, 1994; Goto *et al*, 1997; Ichikawa *et al*, 2001; Harris *et al*, 2010; Bettencourt *et al*, 2010), the extent and regulation of AS, the relative abundance of alternative transcripts in health and disease conditions, the existence of transcript tissue/cell-specificity, and the impact of AS in SCA3/MJD neurotoxicity remain poorly understood. To the best of our knowledge, at least 24 *ATXN3* transcripts, likely to be translated into different isoforms of ataxin-3, have been detected in the human brain and/or peripheral tissues including the blood (Kawaguchi *et al*, 1994; Goto *et al*, 1997; Ichikawa *et al*, 2001; Harris *et al*, 2010; Bettencourt *et al*, 2010), although validation of a large number of such transcripts has not been conducted. *In silico* analysis performed to infer the functional impact of the several putative ataxin-3 protein isoforms showed that some variants were predicted to be “protective” (lacking the CAG tract), whereas others were foreseen to show increased toxicity (Bettencourt *et al*, 2010). Among the *ATXN3* transcripts which are predicted to be translated, only two transcripts encoding ataxin-3 isoforms that differ on the C-terminal containing either two (2UIM) or three UIMs (3UIM) have been mostly studied so far (Harris *et al*, 2010; Johnson *et al*, 2019; Weishäupl *et al*, 2019).

These studies using cellular, mouse or *Drosophila* models and *post-mortem* human brain tissues from healthy individuals, though controversial, uncovered differences in the toxicity of these two ataxin-3 protein isoforms by mainly revealing a faster degradation rate but higher aggregation propensity of the 2UIM isoform compared to the 3UIM (Harris *et al*, 2010; Johnson *et al*, 2019; Weishäupl *et al*, 2019). The fact that these two ataxin-3 isoforms seem to differentially impact SCA3/MJD pathogenesis, raises the possibility that other species generated by *ATXN3* AS play key roles in promoting the selective neuronal toxicity displayed by SCA3/MJD patients.

Several phase I and II clinical trials are ongoing for polyQ disorders aiming to reduce the levels of disease-related transcripts/proteins using ASO-based drugs, but none was yet been approved for clinical use (reviewed in (Helm *et al*, 2022)). Non-allele-specific as well as allele-specific ASO-mediated reduction of *ATXN3* have been successfully tested *in vivo* and *in vitro* models of MJD (McLoughlin *et al*, 2018; Moore *et al*, 2019, 2017; Hauser *et al*, 2022).

Noteworthy, the pharmacokinetics and safety study of a non-selective ASO for SCA3/MJD is currently ongoing (NCT05160558). The identification of specific *ATXN3* transcripts, whose expression needs to be blocked is, therefore, paramount to unveil better strategies in future ASO-based therapies of SCA3/MJD.

Hence, the goal of this study was to profile annotated *ATXN3* AS transcripts in blood and cerebellum of mutation carriers of SCA3/MJD and healthy individuals using RNA sequencing (RNA-seq) data.

## MATERIAL AND METHODS

### Samples and RNA-seq datasets

Two RNA-seq datasets, generated from a total of 72 samples, namely 60 samples from whole blood and 12 samples from *post-mortem* cerebellum (cerebellar lobules) were used in this study: i) a RNA-seq dataset from whole blood samples of 40 SCA3/MJD subjects (30 patients and 10 pre-ataxic) and 20 control individuals (Raposo *et al*, 2023), obtained in the scope of the European spinocerebellar ataxia type 3/Machado-Joseph Initiative (ESMI); and ii) RNA-seq dataset from *post-mortem* cerebellar samples of six SCA3/MJD patients and six controls publicly available from the work of Haas and colleagues (Haas *et al*, 2022). Detailed information about the blood and cerebellum samples from SCA3/MJD subjects (and corresponding healthy controls), is provided in supplementary Table 1. Briefly, RNA extraction and cDNA library preparation was performed according to manufacturer’s instructions, as described elsewhere (Haas *et al*, 2022; Raposo *et al*, 2023). Libraries from blood samples were sequenced as paired end 100bp reads on an Illumina NovaSeq6000 (Illumina), whereas libraries from cerebellum samples were sequenced as paired-end 68 bp reads on a HiSeq2000 (Illumina), both next-generation sequencing (NGS) platforms with limited read length (short-reads).

Ataxin-3 protein levels in plasma (n=28) and in cerebrospinal fluid (CSF, n=8) from a subset of the total 72 SCA3/MJD subjects were available from previous studies(Gonsior *et al*, 2021; Hübener-Schmid *et al*, 2021). Briefly, total full-length ataxin-3 and/or mutant ataxin-3 were measured using a time-resolved fluorescence energy transfer (TR-FRET)-based immunoassay (Gonsior *et al*, 2021) and a single molecule counting (SMC) immunoassay (Hübener-Schmid *et al*, 2021).

The study was approved by local ethics committees of all participating centers; all subjects provided written informed consent.

### Bioinformatic analyses of *ATXN3* splicing

The reads were trimmed using Cutadapt v2.3. Next, reads were mapped to the reference genome GRCh38 (Ensembl), using STAR v2.7.8a (Dobin *et al*, 2013) and assigned count estimates to genes with RSEM v1.3.3(Li & Dewey, 2011). Alignment options followed ENCODE standards for RNA-seq (Dobin *et al*, 2013). FastQC (v0.11.8) (Andrews, 2010) was run on .bam files in a post-alignment step, including both aligned and un-aligned reads, to ensure data quality. From the raw reads, TPM was calculated according to Zhao and colleagues (Zhao *et al*, 2020), considering the denominator as the sum over all samples (brain and blood).

### Ensembl classification biotype

*ATXN3* transcripts were clustered according to the Ensembl classification biotype (Ensembl release 108 – Jan 2023 (Martin *et al*, 2023)) as: (i) protein coding (a transcript that contains an open reading frame), (ii) nonsense mediated decay (NMD; a transcript with a premature stop codon, predicted as most likely to be subjected to targeted degradation because the in-frame termination codon is found more than 50 bp upstream of the final splice junction), (iii) protein coding sequence (CDS) not defined (a transcript of a protein coding gene for which a CDS has not yet been defined), and (iv) retained intron (a transcript that lacks an open reading frame believed to contain intronic sequence relative to other, coding, transcripts of the same gene) which is included in the major category of long non-coding RNA transcripts. Additionally, using the Ensembl annotation for the *ATXN3* gene (Martin *et al*, 2023), *ATXN3* transcripts were clustered according to the presence or absence of the CAG tract, inferred from the presence or absence of exon 10 where the CAG repeat is located.

### Statistical analyses

To increase the sample size and the power of statistical tests, *ATXN3* transcripts were labelled according to its frequency by sample. Transcripts not detected in at least half of our set of samples (i.e., 5 or less samples from cerebellum and 29 or less samples for blood) were not included in statistical analyses (Suppl. Table 2, Fig. S1). Transcripts levels were compared between biological groups (pre-ataxic *versus* controls and patients *versus* controls) and between tissues (cerebellum *versus* blood), and statistical differences were determined by the Mann-Whitney U test. Among the blood samples from controls (n=20), a subset (n=10) was paired-matched by age and sex to pre-ataxic (CTRL-PA) and used for group comparisons (pre-ataxic *versus* CTRL-PA). The significance of the association between absence/presence of transcripts within each tissue by biological group (contingency table) was tested using a Fisher’s exact test. In SCA3/MJD subjects’ group, the relationship between transcript and protein levels was determined using a Spearman rank correlation test. The ROUT method (Q=1%) was used to exclude outliers previously to correlation test. Statistical analyses were performed in GraphPad Prism version 8.0.1 for Windows (GraphPad Software, San Diego, California USA). The significance level of all tests was set to 5%. Graphic bars are shown as median ± 95% CI (confidence interval).

### Availability of data and materials

Data supporting the findings of this study are available in this manuscript and in supplementary material.

## RESULTS

### The abundance and the frequency of annotated *ATXN3* transcripts are higher in cerebellum than in blood regardless of the disease status

Data from RNA-seq of whole blood (n=60) and *post-mortem* cerebellum (n=12) from SCA3/MJD subjects and age- and sex-matched controls was used to identify the currently annotated 54 *ATXN3* transcripts and quantify their abundance (Fig. 1). All annotated transcripts were identified in our study. According to Ensembl annotation (Martin *et al*, 2023), these 54 transcripts can be classified into four main biotypes: protein-coding (n=16), nonsense mediated decay (NMD; n=19), protein CDS not defined (n=16) and retained intron (long noncoding) (n=3) transcripts (Fig. 1). In the 16 protein-coding transcripts group, some structural-related particularities are highlighted (Fig. 1): four transcripts encode full-length ataxin-3 protein isoforms (*ATXN3-*214, *ATXN3-*215, *ATXN3-*227, *ATXN3-*251), including an intact Josephin domain (JD), whereas the remaining 12 transcripts might encode proteins with variations of the N-terminus (*ATXN3*-201, *ATXN3*-203, *ATXN3*-204, *ATXN3*-206, *ATXN3*-207, *ATXN3*-209, *ATXN3*-213, *ATXN3*-219, *ATXN3*-228, *ATXN3*-231, *ATXN3*-235, *ATXN3*-246). Nine transcripts lack exon 11 and, therefore, translate ataxin-3 isoforms only with 2UIM (*ATXN3*- 209, *ATXN3*-213, *ATXN3*-214, *ATXN3*-219, *ATXN3*-227, *ATXN3*-228, *ATXN3*-231, *ATXN3*-235, *ATXN3*-246), while seven transcripts retain exon 11 (*ATXN3*-201, *ATXN3*-203, *ATXN3*-204, *ATXN3*-206, *ATXN3*-207, *ATXN3*-215, *ATXN3*-251), and thus encode 3UIMs isoforms. Finally, and although the annotation of the 3’ untranslated (UTR) is incomplete, two *ATXN3* transcripts seem to lack exon 10, which contains the polyQ-encoding tract (*ATXN3*-209, *ATXN3*-213) (Fig. 1).

**Figure 1.**
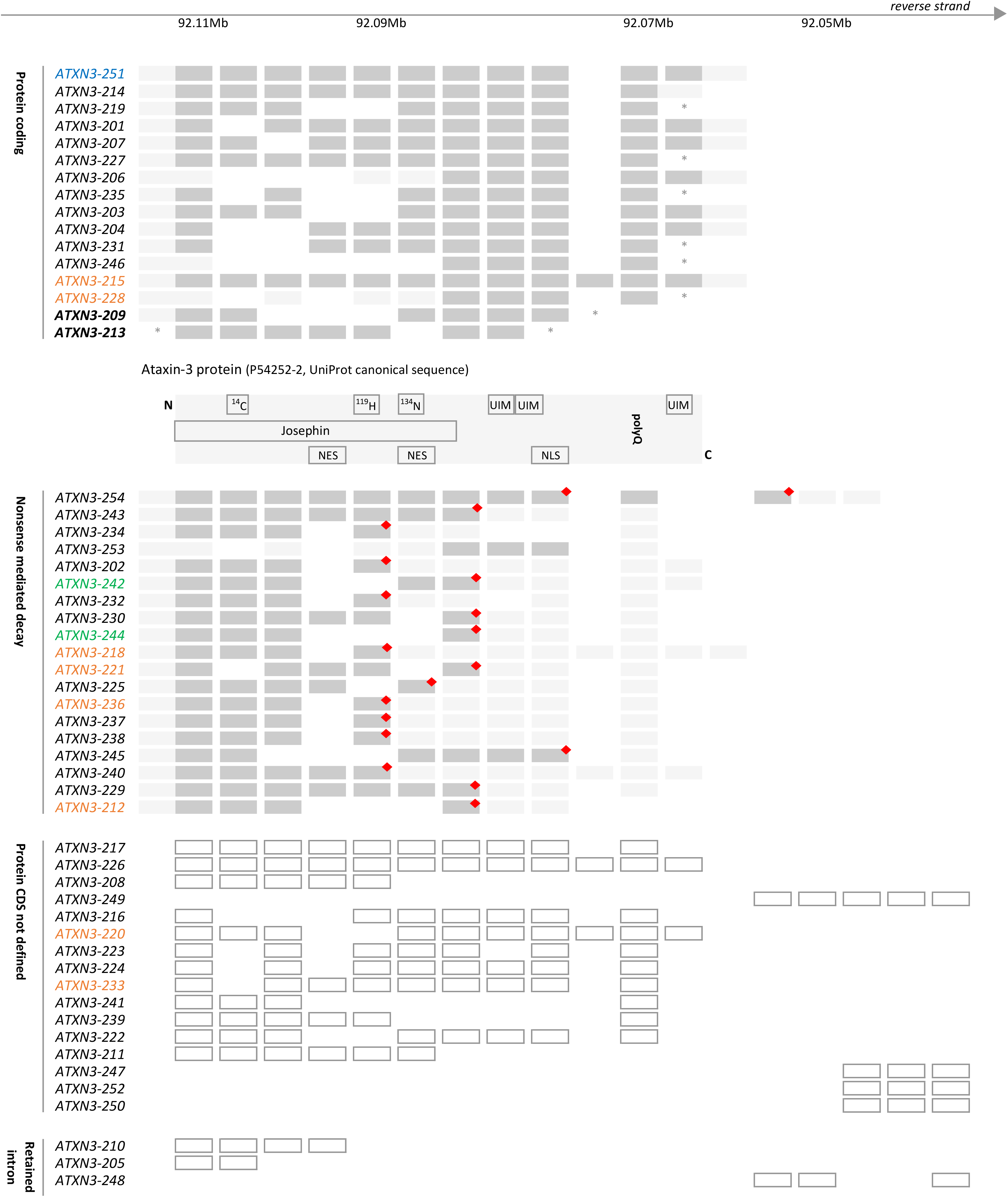
Schematic representation of the annotated 54 *ATXN3* transcripts (adapted from Ensembl release 108 – Jan 2023) detected in blood and cerebellum of pooled SCA3/MJD patients and controls. *ATXN3* transcripts in both SCA3/MJD subjects and controls are clustered according to Ensembl-related biotype - protein coding, nonsense mediated decay, protein coding sequence (CDS) not defined, and retained intron. Moreover, transcripts are clustered by descending order of frequency within tissue calculated in the present study as well as transcripts with exclusive expression in blood and transcripts with exclusive expression in cerebellum are highlighted in orange and green, respectively. The Ensembl’s *ATXN3* reference transcript ENST00000644486.2|*ATXN3*-251 (MANE Select v0.95) is highlighted in blue. Protein-coding transcripts not containing exon 10 (which contains the CAG tract) are highlighted in bold. *The annotation of the transcript is incomplete at 5’ untranslated (UTR), 3’UTR or both. The catalytic amino acids cysteine (C) 14, histidine (H) 119 and asparagine (N) 134 are shown in the respective exons; red diamonds represent a premature termination codon (PTC). Grey box = coding exon; light grey box = UTR; white box = non-coding exon; UIM = ubiquitin interacting motif; polyQ = polyglutamine tract; NES = nuclear export signal; NLS = nuclear localization signal. The size of exons/introns/UTR are not drawn to scale.

To explore putative tissue-specific patterns of *ATXN3* AS regardless of the disease status, the frequency of each annotated *ATXN3* transcript detected in samples from SCA3/MJD subjects and controls was determined within each tissue (blood or cerebellum). Expression levels of *ATXN3* transcripts ranged from 0.01 (blood) to 50.49 (cerebellum) transcripts per million (TPM). Globally, cerebellum samples displayed higher expression levels than blood samples (Fig. 2; Suppl. Table 2). The biotype with more abundant transcripts in blood and cerebellum is the retained intron and the noncoding, respectively (Fig. 2). Thirty out of the 46 (74%) transcripts identified in the cerebellum and 44 out of the 52 (85%) transcripts expressed in the blood could only be detected in less than half of samples (Fig. S1; Suppl. Table 2); for example, the protein-coding transcript *ATXN3*-204 was only expressed in one out of 12 cerebellum samples (8%) and in six out of 60 blood samples (10%) (Suppl. Table 2).

**Figure 2.**
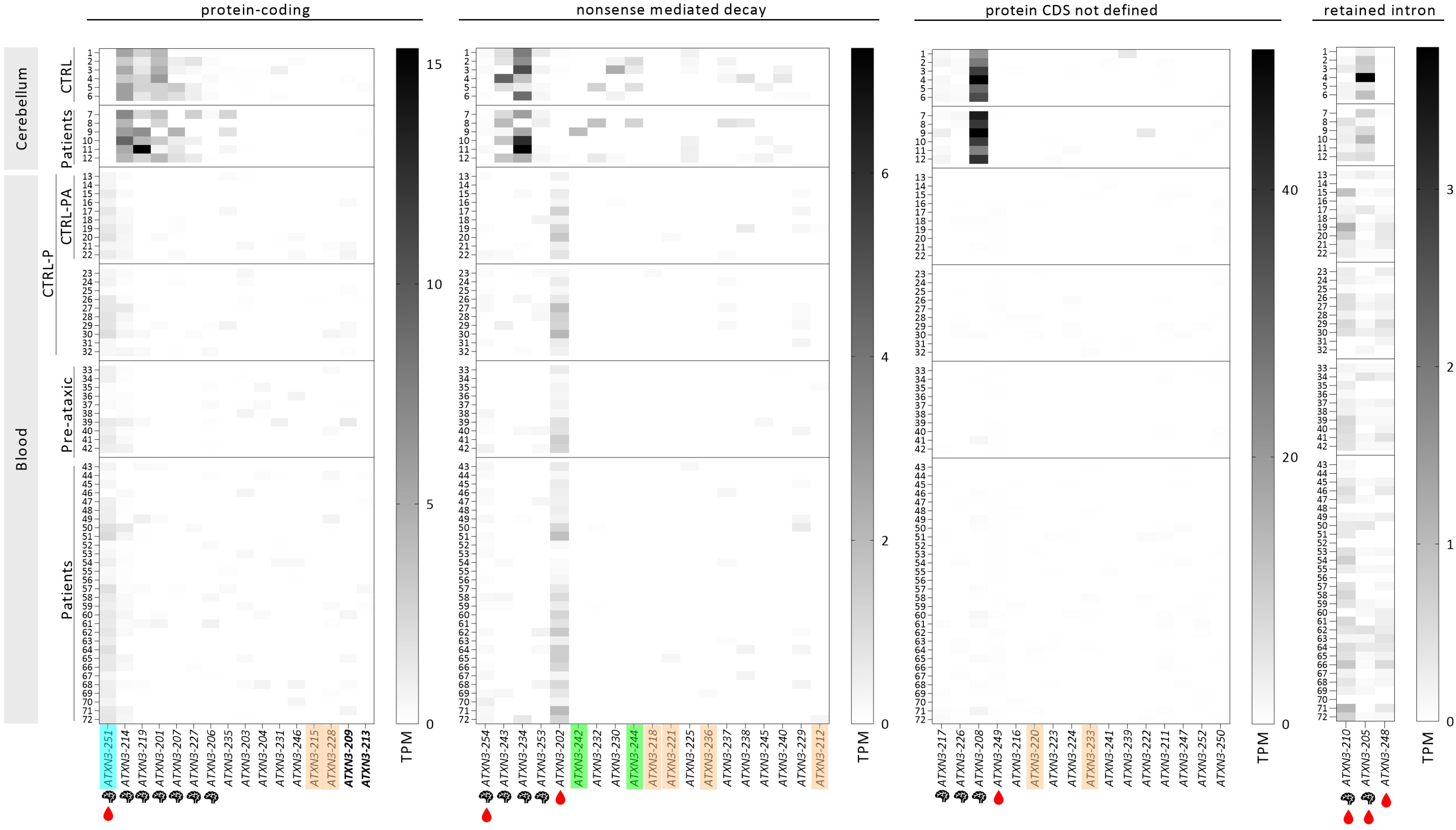
Heatmap of the 54 annotated *ATXN3* transcripts detected in cerebellum and blood samples of SCA3/MJD subjects and healthy individuals. Transcripts are clustered by biotype (protein coding, nonsense mediated decay, protein sequence (CDS) not defined, and retained intron) and disease/health status. Expression values, shown in transcripts per million (TPM), were obtained for 72 samples (12 from cerebellum - lines one to 12) and 60 from whole blood (lines 13 to 72). The magnitude of the expression values was set independently for each transcript type cluster. Protein-coding transcripts not containing exon 10 (which contains the CAG tract) are highlighted in bold. The brain ( ) and/or blood ( ) symbols indicate transcripts detected in more than 50% of all samples in each tissue. *ATXN3* transcripts with exclusive expression in blood and transcripts with exclusive expression in cerebellum are highlighted in orange and green, respectively. The Ensembl’s *ATXN3* reference transcript ENST00000644486.2|*ATXN3*-251 (MANE Select v0.95) is highlighted in blue.

The protein coding *ATXN3*-251 reference transcript and the protein CDS not defined *ATXN3*- 208 transcript showed, respectively, the highest expression in blood and in cerebellum.

Additionally, the transcripts more abundant in blood by biotype were the protein coding *ATXN3*-251, the NMD *ATXN3*-202, the noncoding *ATXN3*-208 and the retained intron *ATXN3*- 210, while in cerebellum the more abundant were the protein coding *ATXN3*-214, the NMD *ATXN3*-234, the noncoding *ATXN3*-208 and the retained intron *ATXN3*-205 (Fig. 2).

Eight of the overall 54 annotated transcripts were exclusively expressed in blood: (i) two corresponded to protein-coding transcripts (*ATXN3*-215, which is one of the four transcripts encoding the full-length ataxin-3 protein isoforms, and *ATXN3*-228); (ii) four were transcripts targeted for NMD (*ATXN3*-212, *ATXN3*-218, *ATXN3*-221, *ATXN3*-236) and (iii) two were protein CDS not defined (*ATXN3*-220, *ATXN3*-233). *ATXN3*-212 transcript showed the highest expression among all transcripts detected in the blood; (Fig. 2, Suppl. Table 2, and Fig. S1). Two NMD transcripts were exclusively expressed in the cerebellum (*ATXN3*-242, *ATXN3*-244) (Suppl. Table 2, Fig. S1).

### Cerebellar abundance of *ATXN3* transcripts is similar in SCA3 patients and controls

Analysis of the distribution of *ATXN3* transcripts by biological group (SCA3/MJD patients and/or controls) in cerebellum samples revealed that several transcripts were detected only in individual or in a limited number of samples (Suppl. Table 2).

To compare distributions of cerebellar *ATXN3* transcripts between SCA3/MJD patients and controls, transcripts detected in more than 50% of all cerebellum samples (six or more of 12 samples) were selected. Following this criterium, 16 transcripts were identified in cerebellum samples corresponding to seven protein-coding (the reference variant - *ATXN3*-251, plus *ATXN3*-201, *ATXN3*-206, *ATXN3*-207, *ATXN3*-214, *ATXN3*-219, *ATXN3*-227), four targeted for NMD (*ATXN3*-234, *ATXN3*-243, *ATXN3*-253, *ATXN3*-254), three protein CDS not defined (*ATXN3*-208, *ATXN3*-217, *ATXN3*-226) and two retained intron (*ATXN3*-205, *ATXN3*-210; Fig. 2).

While all 16 transcripts showed similar abundance in the cerebellum of SCA3/MJD patients and controls (p>0.05; Fig. 3), the protein CDS not defined transcript *ATXN3*-226 was expressed in all samples from controls but was not expressed in four out of the six patients (Fisher’s exact test, p=0.06; Suppl. Table 2).

**Figure 3.**
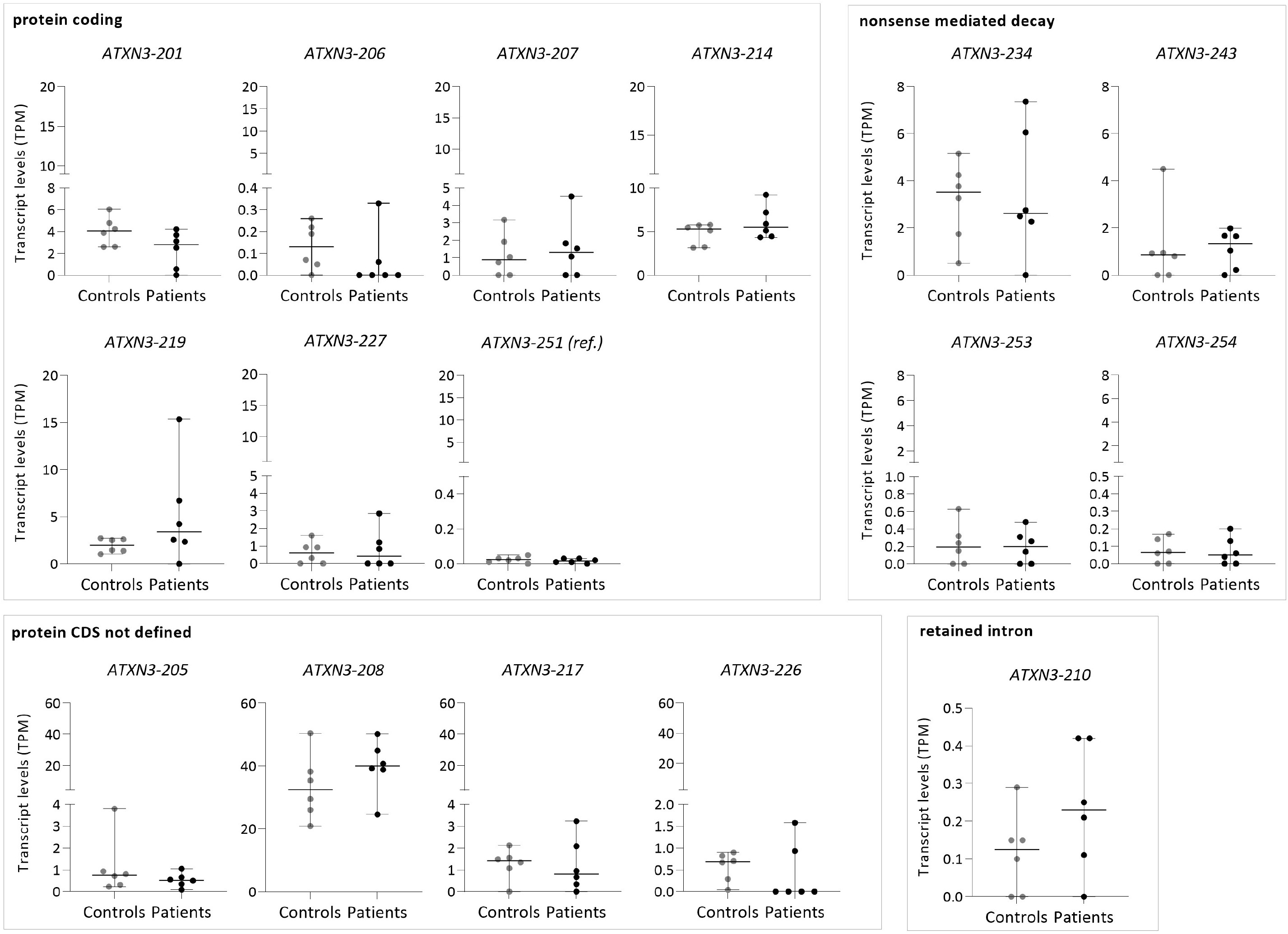
Levels of the 16 *ATXN3* transcripts expressed in more than half of all cerebellum samples from SCA3/MJD patients (n=6) and controls (n=6). The transcripts are clustered according to the type of transcript, as defined by Ensembl classification biotype (Ensembl release 108 - Jan 2023, REF). Transcript expression levels are shown in transcript per million (TPM). Bars in the graphs represent the median ± 95% CI (confidence interval, error bars).

### Blood abundance of *ATXN3* transcripts is similar in SCA3/MJD subjects and controls

Analysis of the distribution of *ATXN3* transcripts in blood samples from SCA3/MJD subjects (pre-ataxic and patients) and controls revealed that, similarly to the cerebellum, several transcripts were only detected in individual or in a limited number of samples (Suppl. Table 2). Following the same criterium used for the cerebellum, eight *ATXN3* transcripts detected in more than 50% of all blood samples (SCA3/MJD subjects and controls) were selected for comparative analyses of patients versus controls and pre-ataxic carriers versus controls. These eight transcripts corresponded to two protein coding (*ATXN3*-214 and the reference transcript - *ATXN3*-251), two transcripts targeted for NMD (*ATXN3*-202, *ATXN3*-254), one protein CDS not defined (*ATXN3*-249), and three retained introns (*ATXN3*-210, *ATXN3*-205, *ATXN3*-248) (Fig. 2). All eight transcripts showed similar levels between SCA3/MJD patients or pre-ataxic carriers compared to their respective controls (Fig. 4).

**Figure 4.**
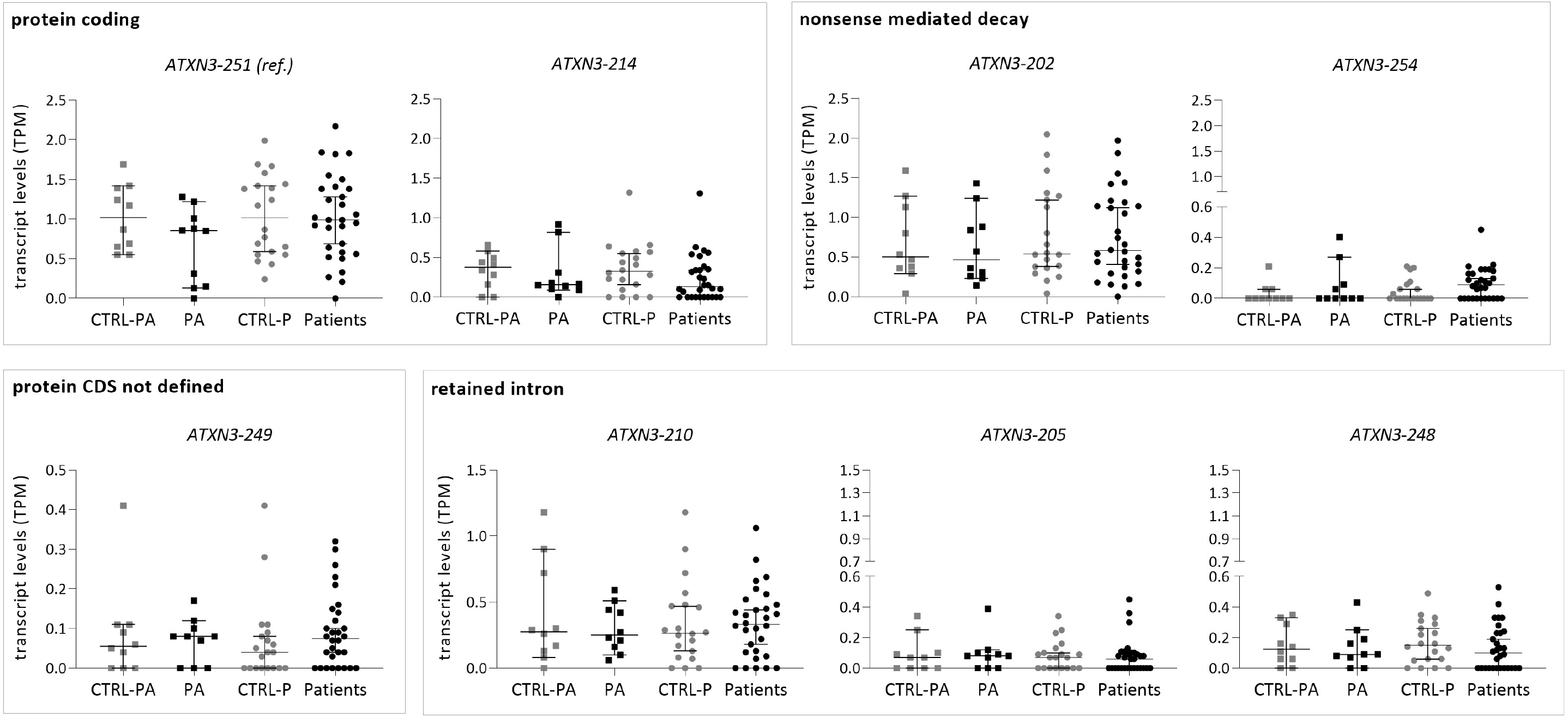
Expression levels of eight *ATXN3* transcripts in blood samples from pre-ataxic (PA) carriers (n=10), SCA3/MJD patients (n=30) and respective controls (CTRL-P, n=20 and CTRL-PA, n=10). CTRL-PA was defined as a subset of age-matched controls with pre-ataxic carriers within the CTRL-P group. The transcripts are clustered according to biotype, as defined by Ensembl classification biotype (Ensembl release 108 - Jan 2023(Martin *et al*, 2023)). Expression levels are shown in transcript per millions (TPM). Bars in the graphs represent the median ± 95% CI (confidence interval, error bars).

### The amount of *ATXN3*-205 transcript in blood is directly associated with CSF levels of soluble mutant ataxin-3

The relationship between *ATXN3* transcript levels from blood cells and soluble levels of ataxin- 3 protein (mutant and/or total) quantified in plasma, peripheral mononuclear blood cells (PBMCs) and CSF samples from the same SCA3/MJD subjects was explored in this study (Fig. S2a). Interestingly, levels of the retained intron transcript *ATXN3*-205 in blood cells directly correlated with the abundance of soluble mutant ataxin-3 in the CSF of SCA3/MJD subjects (p=0.031, Fig. S2c). Although not reaching statistical significance, high levels of *ATXN3*-248 and of the reference transcript *ATXN3*-251 displayed a trend to subtly associate with low levels of total ataxin-3 measured in PBMCs of SCA3/MJD individuals (Fig. S2b, d).

### Expression of the four transcripts encoding full-length ataxin-3 2UIM and 3UIM protein isoforms are tissue-specific, irrespective of disease status

From the 16 protein-coding transcripts, only four encode a full-length ataxin-3 protein with an intact JD and a preserved C-terminus (*ATXN3*-214, *ATXN3*-215, *ATXN3*-227 and *ATXN3*-251; Fig. 5a). *ATXN3*-214 and *ATXN3*-227 transcripts encode ataxin-3 2UIM isoforms, whereas *ATXN3*- 215 and *ATXN3*-251 transcripts encode for ataxin-3 3UIM isoforms (Fig. 5a). To elucidate whether the individual expression levels of the four transcripts encoding full-length ataxin-3 2UIM and 3UIM isoforms contribute to the tissue-specific vulnerability observed in SCA3/MJD, the abundance of these transcripts in blood and cerebellum was compared between SCA3/MJD patients and controls (Fig. 5b).

**Figure 5.**
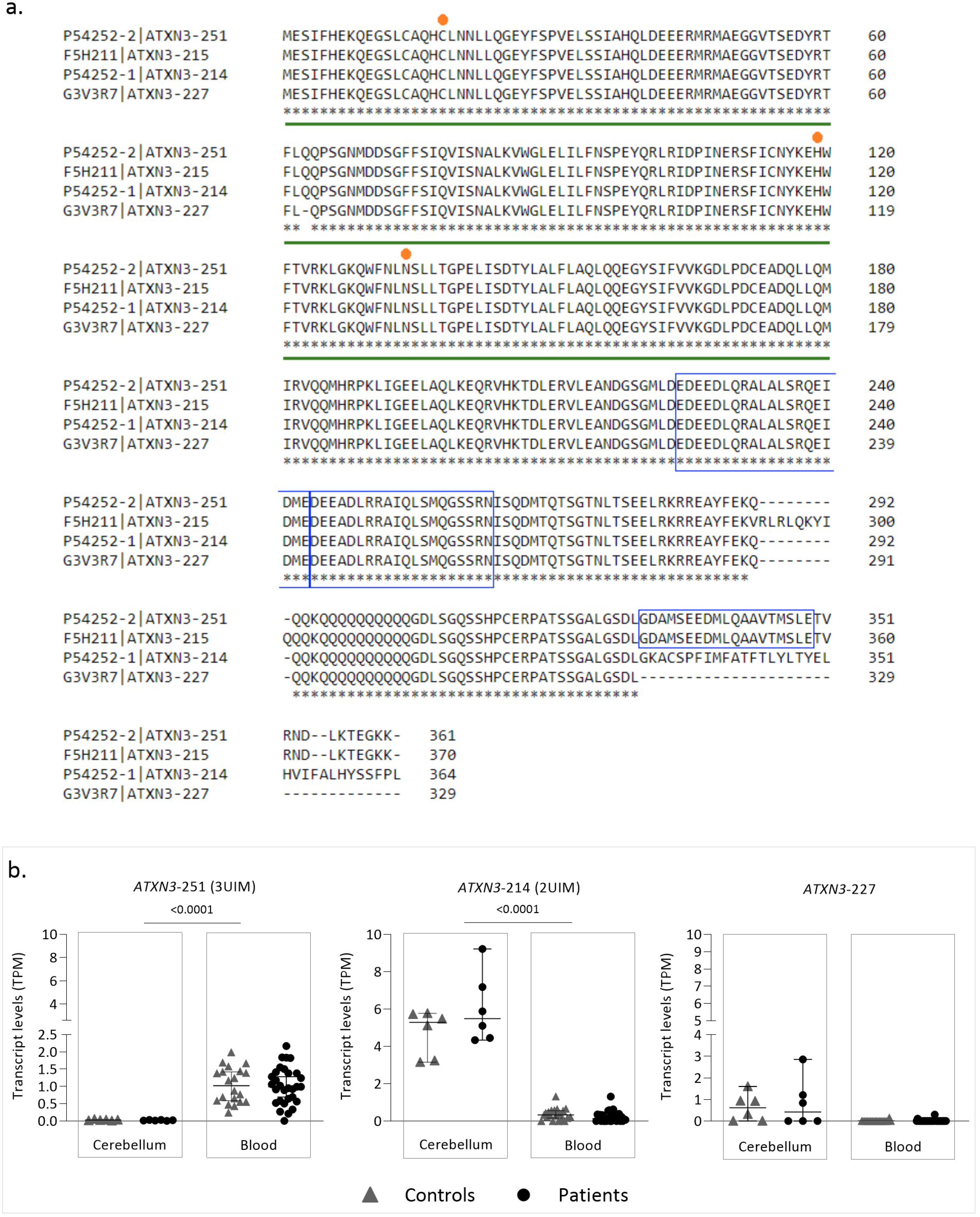
(a) Sequence alignment of the four full-length ataxin-3 protein isoforms harbouring an intact Josephin domain (JD) and a preserved C-terminus. Ataxin-3 protein isoforms, namely the P54252-2, F5H211, P54252-1 and G3V3R7 are encoded by the *ATXN3*-251, *ATXN3*-215, *ATXN3*-214 and *ATXN3*-227 transcripts, respectively. Multiple sequence alignment was performed using Clustal Omega program (Sievers F. et al., 2011). JD (green line), the three catalytic sites (orange dot) and the three UIMs (blue boxes) are highlighted. (b) Levels of *ATXN3*-251 (3UIM) and of the *ATXN3*-214 as well as *ATXN3*-227 (both 2UIM) transcripts in the blood and cerebellum samples from SCA3/MJD subjects and controls. Transcript levels are shown in transcripts per million (TPM). Bars in the graphs represent the median ± 95% CI (confidence interval, error bars).

Interestingly, two of these four transcripts were only expressed in some samples. The *ATXN3*- 215 transcript (3UIM) was only identified in one blood sample (Supp. Table 2); while the *ATXN3*-227 transcript (2UIM) was detected at distinct frequencies in both tissues, it was only measured in three out of 60 (5%) blood samples and in seven out of 12 (58%) in cerebellum samples (Supp. Table 2; Fig. 3). Although being overall commonly detected in the cerebellum, *ATXN3*-227 transcript showed similar levels and frequency in SCA3/MJD patients and controls (Fig. 3).

The reference transcript, *ATXN3*-251 (3UIM), showed a ∼50-fold increase of abundance in blood samples over cerebellum samples, whereas the *ATXN3*-214 (2UIM) transcript was more expressed in the cerebellum about 20-fold more than in blood (Fig. 5b). The magnitude of the fold change values, however, could be partially reflecting technical differences due to different RNA-seq experimental setups. Remarkably, in cerebellum samples, the *ATXN3*-214 (2UIM) was expressed 265-fold more than the *ATXN3*-251 (3UIM) transcript, whereas in blood samples the *ATXN3*-251 (3UIM) transcript was expressed 4-fold more than *ATXN3*-214 (2UIM). These observed differences in transcript abundances between tissues were similar in SCA3/MJD subjects and controls.

## DISCUSSION

Here, we describe the diversity and abundance of annotated *ATXN3* transcripts detected in blood and cerebellum samples from SCA3/MJD subjects and controls. Globally, the number of different transcripts and the abundance of transcripts were higher in the cerebellum than in blood, both in SCA3/MJD subjects and controls. Interestingly, in SCA3/MJD subjects and controls pooled samples altogether, while the most abundant transcript in the cerebellum (*ATXN3*-208) is categorized as a “protein coding with a CDS not defined of unknown function transcript” by Ensembl, the transcript showing the highest abundance in the blood is the protein-coding *ATXN3* reference transcript (*ATXN3*-251) that is translated into an ataxin-3 3UIM protein isoform. No differences in the abundance of the most frequent transcripts were found in blood or cerebellum samples of SCA3/MJD subjects and controls. Noteworthy, the abundance of *ATXN3*-251 and *ATXN3*-214 transcripts, two out of the four transcripts that encode full-length ataxin-3 protein isoforms but differ in the C-terminus (2UIM *versus* 3UIM) were strongly associated with tissue expression specificity: *ATXN3-251* (3UIM) transcript was expressed in blood 50-fold more than in cerebellum, whereas *ATXN3*-214 (2UIM) transcript was expressed in the cerebellum 20-fold more than in blood.

Studies using a *Drosophila* model of SCA3/MJD indicate that in addition to a gain-of-function of the mutant ataxin-3 protein, *ATXN3* transcripts harbouring expanded CAG repeats show independent and progressive cellular toxicity (Li *et al*, 2008). However, whether such toxic transcript action occurs in human cells remains to be elucidated. Hence, the characterization of *ATXN3* AS in mutation carriers of SCA3/MJD might provide fundamental knowledge to unveil the contribution of such mechanism in SCA3/MJD pathogenesis.

Globally, AS shows its highest complexity (high diversity of transcripts), in the brain amongst other tissues (Porter *et al*, 2018). As expected, *ATXN3* transcripts diversity and abundance (a greater number of transcripts, which are expressed in a larger number of samples with the highest levels of expression) was higher in the cerebellum than in the blood and might indicate an important role of *ATXN3* AS and its regulation in the neurodegenerative process of SCA3/MJD.

Twelve out of 16 protein-coding transcripts are predicted to encode proteins with truncations in the JD. Such modifications imply that several ataxin-3 protein forms with a variable N-terminus might exist, displaying for example different forms of the globular JD or other N-terminal domains that show cysteine protease activity or even another yet unknown activity. The fact that ataxin-3 interacts with so many proteins in cells (reviewed in (Costa & Paulson, 2012)) and is ubiquitously expressed (*e.g.,* (Paulson *et al*, 1997)) together with our data indicates that several ataxin-3 isoforms might exist with distinct functions other than the currently known deubiquitinating enzyme role. Interestingly, four out of these 12 transcripts (*ATXN3*-201, *ATXN3*-206, *ATXN3*-207, *ATXN3*-219) are highly expressed in the cerebellum compared to blood suggesting that these potential encoded proteins may play specific roles in the cerebellum and that the correspondent mutant forms could potentially contribute for the observed selective cerebellar toxicity observed in SCA3/MJD patients.

Four out of 54 transcripts, *ATXN3*-214, *ATXN3*-215, *ATXN3*-227 and *ATXN3*-251, encode full-length ataxin-3 protein isoforms, which differ only in the C-terminus. In the present study, the *ATXN3*-214 and the *ATXN3*-227 transcripts, both encoding 2UIM isoforms, are preferentially abundant in cerebellum samples, whereas the *ATXN3*-251 transcript (3UIM) is more expressed in blood. Altogether, these findings suggest that ataxin-3 isoforms with 2UIM might have a unique role in the cerebellum and consequently in neurodegeneration. The correlation between transcript and protein abundance is still not known and future studies exploring the role of the *ATXN3*-214 and *ATXN3-*227 transcripts and respective protein isoforms in brain tissues could contribute for the elucidation of mechanisms involved in cellular vulnerability to mutant *ATXN3* gene products in SCA3/MJD.

Of note, a significant direct relationship between the abundance of the retained intron *ATXN3*- 205 transcript in blood and soluble mutant ataxin-3 in CSF samples was observed in a small sub-set of SCA3/MJD subjects. Because *ATXN3-*205 is classified by Ensembl as a long non-coding RNA, such finding suggests that this transcript might have a regulatory function in the transcription and/or translation of protein coding transcripts in the nervous system and, therefore, contribute to the observed correlation with the soluble mutant ataxin-3 protein in the CSF. More comprehensive studies about the function of these type of transcripts are needed as well as larger number of CSF samples should be further analyzed to better explore this preliminary but important finding.

In this study, transcripts more frequently found in blood (n=8) and in cerebellum (cerebellar lobules, n=16) showed similar levels in SCA3/MJD patients and controls. Mort and colleagues also described similar amount of two *HTT* transcripts in cerebellum samples of HD patients and controls (Mort *et al*, 2015). In SCA3/MJD, neuronal loss in the cerebellar lobules is not as severe as, for example, in the dentate nucleus or pons (Koeppen, 2018). Whether our findings are specific of this brain area is unknown and, thus, characterization of *ATXN3* transcripts in other SCA3/MJD patients brain tissues showing more severe lesions would be important to conduct in the future.

In this work, the diversity and abundance of annotated *ATXN3* transcripts in SCA3/MJD patients and controls is provided. The methodology used in this study did not allow to specifically analyze the *ATXN3* transcripts with CAG repeat expansions. Future studies using other NGS approaches, including novel bioinformatic pipelines to correctly detect CAG expansions are needed to elucidate the contribution of the expanded CAG repeat itself in *ATXN3* AS. Moreover, functional studies to define the role of the most promising transcripts found in the cerebellum, for example for its most abundant known non-coding *ATXN3*-208 transcript, would be important to conduct in the future. In fact, the majority of annotated *ATXN3* transcripts might have distinct functions or regulatory properties and its comprehension will reveal novel mechanisms of SCA3/MJD pathogenesis.

We show that *ATXN3* AS is complex and, thus, its elucidation in the SCA3/MJD-related tissues should become an integral part of future design of *ATXN3* transcripts-targeted therapeutics to overcome potential limitations of efficacy of current RNA-based therapies targeting a limited number of *ATXN3* transcripts. Our findings further aid to the clarification of the specific tissue vulnerability observed in SCA3/MJD and, therefore, may help unravelling novel mechanisms involved in SCA3/MJD pathogenesis.

## ACKNOWLEDGEMENTS

The ESMI consortium would like to thank Nina Roy for coordination and managing of the project. We gratefully thank Dr. Aires Raposo for the collaboration on blood collection in Azores islands.

Additional members of the European Spinocerebellar Ataxia type 3/Machado-Joseph Initiative (ESMI) Study Group are listed in Supplementary Material (Table S3) and participated in subject recruitment and acquisition of participants data as well as read and approved the final manuscript.

## AUTHORS’ ROLES

Design and conceptualization of the study: MR, JH-S, RT, MCC, ML; Subject recruitment/Acquisition of participants biomaterials/data: MR, AFF, ARVM, JV, PP, TK, HG-M, PG, MMS, LPA, JI, BPW, JV, JF, TK, NC, JA, LS, OR, ML; Bioinformatic/Statistical analysis of data: MR, RT; Drafting of the manuscript: MR, JH-S, MCC, ML; Revision of the Manuscript: MR, JH-S, RT, AFF, ARVM, JV, PP, TK, HG-M, PG, MMS, LPA, JI, BPW, JV, JF, TK, NC, JA, LS, OR, MCC, ML.

All authors read and approved the final manuscript.

## REFERENCES

1. Andrews S (2010) FastQC: A Quality Control Tool for High Throughput Sequence Data [Online]. (http://www.bioinformatics.babraham.ac.uk/projects/fastqc/) [PREPRINT]

2. Barbaro BA, Lukacsovich T, Agrawal N, Burke J, Bornemann DJ, Purcell JM, Worthge SA, Caricasole A, Weiss A, Song W, et al (2015) Comparative study of naturally occurring huntingtin fragments in Drosophila points to exon 1 as the most pathogenic species in Huntington’s disease. Hum Mol Genet 24: 913–925

3. Bettencourt C, Santos C, Montiel R, Costa M do C, Cruz-Morales P, Santos LR, Simões N, Kay T, Vasconcelos J, Maciel P, et al (2010) Increased transcript diversity: novel splicing variants of Machado–Joseph Disease gene (ATXN3). Neurogenetics 11: 193–202

4. Burnett B, Li F & Pittman RN (2003) The polyglutamine neurodegenerative protein ataxin-3 binds polyubiquitylated proteins and has ubiquitin protease activity. Hum Mol Genet 12: 3195–205

5. Costa M do C & Paulson HL (2012) Toward understanding Machado-Joseph disease. Prog Neurobiol 97: 239–257 doi:10.1016/j.pneurobio.2011.11.006 [PREPRINT]

6. Daguenet E, Dujardin G & Valcárcel J (2015) The pathogenicity of splicing defects: mechanistic insights into pre- mRNA processing inform novel therapeutic approaches. EMBO Rep 16: 1640–1655

7. Dhuri K, Bechtold C, Quijano E, Pham H, Gupta A, Vikram A & Bahal R (2020) Antisense Oligonucleotides: An Emerging Area in Drug Discovery and Development. J Clin Med 9: 2004

8. Dobin A, Davis CA, Schlesinger F, Drenkow J, Zaleski C, Jha S, Batut P, Chaisson M & Gingeras TR (2013) STAR: ultrafast universal RNA-seq aligner. Bioinformatics 29: 15–21

9. Gonsior K, Kaucher GA, Pelz P, Schumann D, Gansel M, Kuhs S, Klockgether T, Forlani S, Durr A, Hauser S, et al (2021) PolyQ-expanded ataxin-3 protein levels in peripheral blood mononuclear cells correlate with clinical parameters in SCA3: a pilot study. J Neurol 268: 1304–1315

10. Goto J, Watanabe M, Ichikawa Y, Yee S-B, Ihara N, Endo K, Igarashi S, Takiyama Y, Gaspar C, Maciel P, et al (1997) Machado–Joseph disease gene products carrying different carboxyl termini. Neurosci Res 28: 373–377

11. Haas E, Incebacak RD, Hentrich T, Huridou C, Schmidt T, Casadei N, Maringer Y, Bahl C, Zimmermann F, Mills JD, et al (2022) A Novel SCA3 Knock-in Mouse Model Mimics the Human SCA3 Disease Phenotype Including Neuropathological, Behavioral, and Transcriptional Abnormalities Especially in Oligodendrocytes. Mol Neurobiol 59: 495–522

12. Harris GM, Dodelzon K, Gong L, Gonzalez-Alegre P & Paulson HL (2010) Splice isoforms of the polyglutamine disease protein ataxin-3 exhibit similar enzymatic yet different aggregation properties. PLoS One 5

13. Hauser S, Helm J, Kraft M, Korneck M, Hübener-Schmid J & Schöls L (2022) Allele-specific targeting of mutant ataxin-3 by antisense oligonucleotides in SCA3-iPSC-derived neurons. Mol Ther - Nucleic Acids 27: 99–108

14. Helm J, Schöls L & Hauser S (2022) Towards Personalized Allele-Specific Antisense Oligonucleotide Therapies for Toxic Gain-of-Function Neurodegenerative Diseases. Pharmaceutics 14: 1708

15. Hübener-Schmid J, Kuhlbrodt K, Peladan J, Faber J, Santana MM, Hengel H, Jacobi H, Reetz K, Garcia-Moreno H, Raposo M, et al (2021) Polyglutamine-Expanded Ataxin-3: A Target Engagement Marker for Spinocerebellar Ataxia Type 3 in Peripheral Blood. Mov Disord 36: 2675–2681

16. Ichikawa Y, Goto J, Hattori M, Toyoda A, Ishii K, Jeong SY, Hashida H, Masuda N, Ogata K, Kasai F, et al (2001) The genomic structure and expression of MJD, the Machado-Joseph disease gene. J Hum Genet 46: 413–22

17. Johnson SL, Blount JR, Libohova K, Ranxhi B, Paulson HL, Tsou W-L & Todi S V. (2019) Differential toxicity of ataxin-3 isoforms in Drosophila models of Spinocerebellar Ataxia Type 3. Neurobiol Dis 132: 104535

18. Kawaguchi Y, Okamoto T, Taniwaki M, Aizawa M, Inoue M, Katayama S, Kawakami H, Nakamura S, Nishimura M & Akiguchi I (1994) CAG expansions in a novel gene for Machado-Joseph disease at chromosome 14q32.1. Nat Genet 8: 221–8

19. Koeppen AH (2018) The Neuropathology of Spinocerebellar Ataxia Type 3/Machado-Joseph Disease. In pp 233–241.

20. Li B & Dewey CN (2011) RSEM: accurate transcript quantification from RNA-Seq data with or without a reference genome. BMC Bioinformatics 12: 323

21. Li L-B, Yu Z, Teng X & Bonini NM (2008) RNA toxicity is a component of ataxin-3 degeneration in Drosophila. Nature 453: 1107–1111

22. Maciel P, Costa MC, Ferro A, Rousseau M, Santos CS, Gaspar C, Barros J, Rouleau GA, Coutinho P & Sequeiros J (2001) Improvement in the molecular diagnosis of Machado-Joseph disease. Arch Neurol 58: 1821–1827

23. Mangiarini L, Sathasivam K, Seller M, Cozens B, Harper A, Hetherington C, Lawton M, Trottier Y, Lehrach H, Davies SW, et al (1996) Exon 1 of the HD Gene with an Expanded CAG Repeat Is Sufficient to Cause a Progressive Neurological Phenotype in Transgenic Mice. Cell 87: 493–506

24. Martin FJ, Amode MR, Aneja A, Austine-Orimoloye O, Azov AG, Barnes I, Becker A, Bennett R, Berry A, Bhai J, et al (2023) Ensembl 2023. Nucleic Acids Res 51: D933–D941

25. Masino L, Musi V, Menon RP, Fusi P, Kelly G, Frenkiel TA, Trottier Y & Pastore A (2003) Domain architecture of the polyglutamine protein ataxin-3: a globular domain followed by a flexible tail. FEBS Lett 549: 21–5

26. McLoughlin HS, Moore LR, Chopra R, Komlo R, McKenzie M, Blumenstein KG, Zhao H, Kordasiewicz HB, Shakkottai VG & Paulson HL (2018) Oligonucleotide therapy mitigates disease in spinocerebellar ataxia type 3 mice. Ann Neurol 84: 64–77

27. Moore LR, Keller L, Bushart DD, Delatorre RG, Li D, McLoughlin HS, do Carmo Costa M, Shakkottai VG, Smith GD & Paulson HL (2019) Antisense oligonucleotide therapy rescues aggresome formation in a novel spinocerebellar ataxia type 3 human embryonic stem cell line. Stem Cell Res 39: 101504

28. Moore LR, Rajpal G, Dillingham IT, Qutob M, Blumenstein KG, Gattis D, Hung G, Kordasiewicz HB, Paulson HL & McLoughlin HS (2017) Evaluation of Antisense Oligonucleotides Targeting ATXN3 in SCA3 Mouse Models. Mol Ther - Nucleic Acids 7: 200–210

29. Mort M, Carlisle FA, Waite AJ, Elliston L, Allen ND, Jones L & Hughes AC (2015) Huntingtin Exists as Multiple Splice Forms in Human Brain. J Huntingtons Dis 4: 161–171

30. Neueder A, Landles C, Ghosh R, Howland D, Myers RH, Faull RLM, Tabrizi SJ & Bates GP (2017) The pathogenic exon 1 HTT protein is produced by incomplete splicing in Huntington’s disease patients. Sci Rep 7: 1307

31. Nik S & Bowman T V. (2019) Splicing and neurodegeneration: Insights and mechanisms. WIREs RNA 10

32. Paulson HL, Das SS, Crino PB, Perez MK, Patel SC, Gotsdiner D, Fischbeck KH & Pittman RN (1997) Machado-Joseph disease gene product is a cytoplasmic protein widely expressed in brain. Ann Neurol 41: 453–62

33. Porter RS, Jaamour F & Iwase S (2018) Neuron-specific alternative splicing of transcriptional machineries: Implications for neurodevelopmental disorders. Mol Cell Neurosci 87: 35–45

34. Raposo M, Hübener-Schmid J, Ferreira AF, Vieira Melo AR, Vasconcelos J, Pires P, Kay T, Garcia- Moreno H, Giunti P, Santana MM, et al (2023) Blood transcriptome sequencing identifies biomarkers able to track disease stages in spinocerebellar ataxia type 3. Brain accepted

35. Sathasivam K, Neueder A, Gipson TA, Landles C, Benjamin AC, Bondulich MK, Smith DL, Faull RLM, Roos RAC, Howland D, et al (2013) Aberrant splicing of HTT generates the pathogenic exon 1 protein in Huntington disease. Proc Natl Acad Sci 110: 2366–2370

36. Sequeiros J, Martins S & Silveira I (2012) Epidemiology and population genetics of degenerative ataxias. In pp 227–251.

37. Takiyama Y, Nishizawa M, Tanaka H, Kawashima S, Sakamoto H, Karube Y, Shimazaki H, Soutome M, Endo K & Ohta S (1993) The gene for Machado-Joseph disease maps to human chromosome 14q. Nat Genet 4: 300–4

38. Vuong CK, Black DL & Zheng S (2016) The neurogenetics of alternative splicing. Nat Rev Neurosci 17: 265–281

39. Weishäupl D, Schneider J, Peixoto Pinheiro B, Ruess C, Dold SM, von Zweydorf F, Gloeckner CJ, Schmidt J, Riess O & Schmidt T (2019) Physiological and pathophysiological characteristics of ataxin-3 isoforms. J Biol Chem 294: 644–661

40. Zhao S, Ye Z & Stanton R (2020) Misuse of RPKM or TPM normalization when comparing across samples and sequencing protocols. RNA 26: 903–909

41. Zhu Y, Zhu L, Wang X & Jin H (2022) RNA-based therapeutics: an overview and prospectus. Cell Death Dis 13: 644

