## Supplementary material for "Blood and cerebellar abundance of *ATXN3* splice variants in spinocerebellar ataxia type 3/Machado-Joseph disease"

Supplementary Table 1. Demographic and genetic information of healthy individuals and carriers of the *ATXN3* mutation (pre-ataxic subjects and patients) included in this study and grouped by type of tissue - whole blood and cerebellum.

Whole blood

| | **CTRL-PA** | **Pre-ataxic carriers (PA)** | **CTRL-P** | **Patients (P)** | Group comparisons |
|---|---|---|---|---|---|
| Sample size, n | 10 | 10 | 20 | 30 | |
| Gender (Female\|Male) | 5\|5 | 5\|5 | 10\|10 | 15\|15 | CTRL-PA = PA; CTRL-P = P; PA = P |
| Age, years | 34 [28-42] | 36 [30-40] | 49 [33-62] | 52 [43-62] | CTRL-PA = PA; CTRL-P = P; PA ≠ P |
| $CAG_n$ allele 1 | 14 [14-20] | 18 [14-23] | 14 [14-23] | 23 [20-26] | CTRL-PA = PA; CTRL-P ≠ P |
| $CAG_n$ allele 2 | 23 [23-25] | 69 [64-71] | 23 [23-27] | 70 [66-72] | PA = P |
| Age at onset (AO), years | - | 35 [32-36], n=4 | - | 38 [26-47] | CTRL-PA = PA; CTRL-P = P; PA = P |
| SARA score | 0 [0-0.1] | 1 [0-1] | 0 [0-0.5] | 18 [6-28] | CTRL-PA ≠ PA; PA ≠ P; CTRL-P ≠ P |

Cerebellum

| | | **CTRL-P** | **Patients (P)** | |
|---|---|---|---|---|
| Sample size, n | | 6 | 6 | |
| Gender (Female\|Male) | | 0\|6 | 0\|6 | CTRL-P = P |
| Age, years | | 71 [51-77] | 68 [58-75] | CTRL-P = P |

Continuous variables are shown as median [Interquartile range:1stQ-3thQ]. For whole blood, among the total number of controls (CTRL-P; n=20), a subset (n=10) was selected to match the pre-ataxic carriers (PA) by age and sex. SARA scores[a] were used to classify SCA3/MJD mutation carriers as either patients (SARA score ≥3) or pre-ataxic subjects (SARA score <3)[b]. Age was calculated as the difference between the year of birth and the year of the clinical evaluation/blood collection. The number of CAG repeats ($CAG_n$) in the two *ATXN3* alleles (allele 1 and allele 2) is shown for all individuals. AO was defined as the age of the first gait disturbances, reported by the patient or a close relative/caregiver. Group comparisons for whole blood and cerebellum samples were conducted using a chi-square test of independence to compare the proportion of subjects by gender, or by Mann-Whitney U test to compare age, the number of CAG repeats in *ATXN3*, AO, and SARA score and by setting the statistical significance as $p < 0.05$ (≠).

a. Schmitz-Hübsch T, du Montcel ST, Baliko L, Berciano J, Boesch S, Depondt C, et al. Scale for the assessment and rating of ataxia: development of a new clinical scale. Neurology. 2006 Jun;66(11):1717–20.
b. Maas RPPWM, van Gaalen J, Klockgether T, van de Warrenburg BPC. The preclinical stage of spinocerebellar ataxias. Neurology. 2015 Jul 7;85(1):96–103.

**Supplementary table 2.** *ATXN3* transcript levels (in TPM, transcripts per million) by sample (n=72) according to type of tissue (a) and determination of absolute and relative frequencies of each splice variant (n=54) by biological group according to type of tissue (b). *ATXN3* transcripts are clustered by biotype (protein coding, nonsense mediated decay, protein coding sequence (CDS) not defined and retained intron transcripts).

**a.**

| | | Protein coding | | | | | | | | | | | | | | | |
|---|---|---|---|---|---|---|---|---|---|---|---|---|---|---|---|---|---|
| | | ATXN3-251 (ref.) | ATXN3-201 | ATXN3-203 | ATXN3-204 | ATXN3-206 | ATXN3-207 | ATXN3-209 | ATXN3-213 | ATXN3-214 | ATXN3-215 | ATXN3-219 | ATXN3-227 | ATXN3-228 | ATXN3-231 | ATXN3-235 | ATXN3-246 |
| Cerebellum | CTRL1 | 0.03 | 4.25 | 0.00 | 0.00 | 0.00 | 0.00 | 0.00 | 0.00 | 5.47 | 0.00 | 2.73 | 0.00 | 0.00 | 0.00 | 0.00 | 0.00 |
| | CTRL2 | 0.05 | 3.89 | 0.00 | 0.00 | 0.22 | 1.04 | 0.00 | 0.00 | 3.15 | 0.00 | 1.04 | 1.60 | 0.00 | 0.00 | 0.00 | 0.19 |
| | CTRL3 | 0.00 | 4.79 | 0.04 | 0.01 | 0.05 | 0.73 | 0.00 | 0.00 | 5.11 | 0.00 | 1.47 | 0.31 | 0.00 | 0.91 | 0.12 | 0.00 |
| | CTRL4 | 0.03 | 6.04 | 0.00 | 0.00 | 0.26 | 0.00 | 0.12 | 0.00 | 3.24 | 0.00 | 2.52 | 0.00 | 0.00 | 0.00 | 0.00 | 0.00 |
| | CTRL5 | 0.02 | 2.60 | 0.01 | 0.00 | 0.07 | 3.18 | 0.00 | 0.22 | 5.77 | 0.00 | 2.60 | 0.92 | 0.00 | 0.00 | 0.00 | 0.00 |
| | CTRL6 | 0.01 | 2.61 | 0.00 | 0.00 | 0.19 | 1.91 | 0.00 | 0.00 | 5.72 | 0.00 | 1.39 | 0.93 | 0.00 | 0.13 | 0.00 | 0.00 |
| | Patient1 | 0.03 | 3.69 | 0.00 | 0.00 | 0.33 | 0.00 | 0.00 | 0.00 | 7.18 | 0.00 | 2.35 | 2.86 | 0.00 | 0.00 | 2.52 | 0.00 |
| | Patient2 | 0.01 | 2.52 | 0.00 | 0.00 | 0.00 | 0.00 | 0.00 | 0.00 | 4.45 | 0.00 | 0.00 | 0.00 | 0.00 | 0.00 | 0.00 | 0.00 |
| | Patient3 | 0.01 | 0.00 | 0.00 | 0.00 | 0.00 | 4.52 | 0.00 | 0.07 | 5.88 | 0.00 | 6.73 | 0.00 | 0.00 | 0.00 | 1.71 | 0.00 |
| | Patient4 | 0.03 | 3.10 | 0.00 | 0.00 | 0.00 | 1.07 | 0.00 | 0.00 | 9.22 | 0.00 | 4.24 | 0.84 | 0.00 | 0.00 | 0.00 | 0.00 |
| | Patient5 | 0.00 | 0.55 | 0.02 | 0.00 | 0.00 | 1.53 | 0.00 | 0.00 | 5.10 | 0.00 | 15.35 | 0.00 | 0.00 | 0.00 | 0.54 | 0.00 |
| | Patient6 | 0.02 | 4.21 | 0.00 | 0.00 | 0.06 | 1.82 | 0.00 | 0.00 | 4.34 | 0.00 | 2.56 | 1.21 | 0.00 | 0.00 | 0.00 | 0.00 |
| Blood | CTRL1 | 0.59 | 0.00 | 0.47 | 0.00 | 0.00 | 0.00 | 0.00 | 0.00 | 0.22 | 0.00 | 0.00 | 0.00 | 0.00 | 0.00 | 0.00 | 0.00 |
| | CTRL2 | 0.77 | 0.00 | 0.22 | 0.00 | 0.00 | 0.08 | 0.00 | 0.00 | 0.09 | 0.00 | 0.00 | 0.00 | 0.00 | 0.00 | 0.00 | 0.00 |
| | CTRL3 | 0.55 | 0.00 | 0.05 | 0.00 | 0.00 | 0.00 | 0.00 | 0.00 | 0.16 | 0.00 | 0.00 | 0.00 | 0.00 | 0.00 | 0.22 | 0.00 |
| | CTRL4 | 0.24 | 0.00 | 0.00 | 0.00 | 0.00 | 0.00 | 0.13 | 0.00 | 0.00 | 0.00 | 0.00 | 0.00 | 0.00 | 0.00 | 0.00 | 0.00 |
| | CTRL5 | 0.55 | 0.00 | 0.00 | 0.00 | 0.00 | 0.00 | 0.00 | 0.00 | 0.00 | 0.00 | 0.00 | 0.00 | 0.00 | 0.00 | 0.00 | 0.00 |
| | CTRL6 | 1.24 | 0.00 | 0.00 | 0.00 | 0.00 | 0.00 | 0.00 | 0.00 | 0.28 | 0.00 | 0.00 | 0.00 | 0.00 | 0.00 | 0.00 | 0.00 |
| | CTRL7 | 1.38 | 0.00 | 0.14 | 0.00 | 0.00 | 0.00 | 0.00 | 0.04 | 0.23 | 0.05 | 0.00 | 0.00 | 0.00 | 0.00 | 0.00 | 0.00 |
| | CTRL8 | 0.69 | 0.00 | 0.00 | 0.00 | 0.00 | 0.00 | 0.27 | 0.00 | 0.00 | 0.00 | 0.00 | 0.00 | 0.00 | 0.00 | 0.00 | 0.00 |
| | CTRL9 | 1.44 | 0.00 | 0.05 | 0.00 | 0.00 | 0.00 | 0.00 | 0.00 | 1.32 | 0.00 | 0.15 | 0.00 | 0.00 | 0.00 | 0.00 | 0.00 |
| | CTRL10 | 1.39 | 0.00 | 0.00 | 0.00 | 0.00 | 0.00 | 0.00 | 0.00 | 0.34 | 0.00 | 0.00 | 0.00 | 0.00 | 0.00 | 0.43 | 0.00 |
| | CTRL11 | 0.87 | 0.00 | 0.00 | 0.00 | 0.00 | 0.18 | 0.00 | 0.00 | 0.41 | 0.00 | 0.00 | 0.00 | 0.00 | 0.00 | 0.00 | 0.00 |
| | CTRL12 | 1.42 | 0.00 | 0.00 | 0.00 | 0.00 | 0.14 | 0.00 | 0.00 | 0.66 | 0.00 | 0.00 | 0.00 | 0.00 | 0.00 | 0.00 | 0.00 |
| | CTRL13 | 1.69 | 0.27 | 0.00 | 0.00 | 0.00 | 0.00 | 0.00 | 0.00 | 0.49 | 0.00 | 0.00 | 0.00 | 0.00 | 0.05 | 0.00 | 0.31 |
| | CTRL14 | 0.65 | 0.00 | 0.48 | 0.00 | 0.00 | 0.00 | 0.38 | 0.00 | 0.44 | 0.00 | 0.00 | 0.00 | 0.18 | 0.00 | 0.00 | 0.00 |
| | CTRL15 | 1.67 | 0.00 | 0.00 | 0.00 | 0.00 | 0.00 | 0.00 | 0.00 | 0.57 | 0.00 | 0.00 | 0.00 | 0.00 | 0.00 | 0.00 | 0.00 |
| | CTRL16 | 1.17 | 0.00 | 0.00 | 0.00 | 0.00 | 0.28 | 0.66 | 0.00 | 0.58 | 0.00 | 0.00 | 0.00 | 0.01 | 0.00 | 0.00 | 0.40 |
| | CTRL17 | 1.58 | 0.24 | 0.00 | 0.00 | 0.12 | 0.00 | 0.32 | 0.00 | 0.31 | 0.00 | 0.00 | 0.00 | 0.00 | 0.00 | 0.69 | 0.00 |
| | CTRL18 | 1.99 | 0.00 | 0.00 | 0.00 | 0.00 | 0.00 | 0.31 | 0.00 | 0.64 | 0.00 | 0.31 | 0.16 | 0.56 | 0.00 | 0.00 | 0.00 |
| | CTRL19 | 0.43 | 0.00 | 0.00 | 0.00 | 0.00 | 0.00 | 0.00 | 0.00 | 0.00 | 0.00 | 0.00 | 0.00 | 0.00 | 0.00 | 0.00 | 0.00 |
| | CTRL20 | 0.47 | 0.00 | 0.00 | 0.00 | 0.36 | 0.05 | 0.09 | 0.00 | 0.55 | 0.00 | 0.32 | 0.00 | 0.00 | 0.00 | 0.00 | 0.00 |
| | Patient1 | 0.64 | 0.20 | 0.00 | 0.00 | 0.00 | 0.00 | 0.00 | 0.00 | 0.00 | 0.00 | 0.29 | 0.00 | 0.00 | 0.00 | 0.00 | 0.00 |
| | Patient2 | 0.21 | 0.00 | 0.00 | 0.22 | 0.08 | 0.00 | 0.00 | 0.09 | 0.00 | 0.00 | 0.00 | 0.00 | 0.40 | 0.05 | 0.00 | 0.00 |
| | Patient3 | 0.58 | 0.00 | 0.00 | 0.00 | 0.00 | 0.00 | 0.00 | 0.00 | 0.00 | 0.00 | 0.00 | 0.00 | 0.02 | 0.00 | 0.00 | 0.00 |
| | Patient4 | 0.00 | 0.00 | 0.63 | 0.00 | 0.00 | 0.00 | 0.00 | 0.00 | 0.34 | 0.00 | 0.00 | 0.00 | 0.00 | 0.00 | 0.00 | 0.00 |
| | Patient5 | 0.89 | 0.00 | 0.00 | 0.00 | 0.00 | 0.00 | 0.00 | 0.10 | 0.11 | 0.00 | 0.00 | 0.00 | 0.00 | 0.00 | 0.00 | 0.00 |
| | Patient6 | 1.02 | 0.00 | 0.00 | 0.00 | 0.00 | 0.00 | 0.00 | 0.00 | 0.09 | 0.00 | 0.00 | 0.00 | 0.00 | 0.00 | 0.00 | 0.00 |
| | Patient7 | 0.99 | 0.29 | 0.00 | 0.00 | 0.00 | 0.00 | 0.00 | 0.00 | 0.00 | 0.00 | 0.85 | 0.00 | 0.67 | 0.00 | 0.00 | 0.00 |
| | Patient8 | 1.84 | 0.00 | 0.00 | 0.00 | 0.00 | 0.00 | 0.00 | 0.00 | 1.31 | 0.00 | 0.00 | 0.30 | 0.01 | 0.00 | 0.00 | 0.00 |
| | Patient9 | 2.17 | 0.00 | 0.00 | 0.00 | 0.00 | 0.00 | 0.00 | 0.00 | 0.56 | 0.00 | 0.03 | 0.00 | 0.00 | 0.00 | 0.00 | 0.00 |
| | Patient10 | 0.27 | 0.00 | 0.00 | 0.00 | 0.20 | 0.00 | 0.00 | 0.00 | 0.00 | 0.00 | 0.00 | 0.00 | 0.00 | 0.01 | 0.00 | 0.00 |
| | Patient11 | 0.52 | 0.00 | 0.41 | 0.00 | 0.00 | 0.00 | 0.00 | 0.00 | 0.07 | 0.00 | 0.00 | 0.00 | 0.00 | 0.00 | 0.00 | 0.00 |
| | Patient12 | 0.95 | 0.00 | 0.00 | 0.00 | 0.00 | 0.00 | 0.00 | 0.00 | 0.15 | 0.00 | 0.00 | 0.00 | 0.00 | 0.41 | 0.00 | 0.42 |
| | Patient13 | 0.33 | 0.00 | 0.00 | 0.00 | 0.00 | 0.00 | 0.00 | 0.00 | 0.10 | 0.00 | 0.00 | 0.00 | 0.00 | 0.00 | 0.17 | 0.00 |
| | Patient14 | 0.50 | 0.00 | 0.00 | 0.00 | 0.00 | 0.00 | 0.00 | 0.00 | 0.10 | 0.00 | 0.00 | 0.00 | 0.00 | 0.00 | 0.00 | 0.03 |
| | Patient15 | 1.82 | 0.00 | 0.00 | 0.00 | 0.00 | 0.08 | 0.00 | 0.37 | 0.15 | 0.00 | 0.22 | 0.00 | 0.00 | 0.00 | 0.31 | 0.00 |
| | Patient16 | 1.18 | 0.00 | 0.00 | 0.00 | 0.48 | 0.00 | 0.00 | 0.00 | 0.32 | 0.00 | 0.00 | 0.00 | 0.00 | 0.00 | 0.00 | 0.00 |
| | Patient17 | 0.92 | 0.00 | 0.30 | 0.00 | 0.00 | 0.00 | 0.00 | 0.12 | 0.35 | 0.00 | 0.00 | 0.00 | 0.22 | 0.00 | 0.00 | 0.00 |
| | Patient18 | 1.38 | 0.18 | 0.00 | 0.21 | 0.00 | 0.00 | 0.49 | 0.00 | 0.54 | 0.00 | 0.00 | 0.00 | 0.00 | 0.00 | 0.00 | 0.00 |
| | Patient19 | 1.24 | 0.66 | 0.00 | 0.00 | 0.74 | 0.00 | 0.00 | 0.00 | 0.25 | 0.00 | 0.39 | 0.00 | 0.10 | 0.00 | 0.00 | 0.00 |
| | Patient20 | 1.50 | 0.00 | 0.00 | 0.00 | 0.00 | 0.00 | 0.00 | 0.00 | 0.34 | 0.00 | 0.00 | 0.00 | 0.00 | 0.00 | 0.00 | 0.00 |
| | Patient21 | 1.06 | 0.00 | 0.00 | 0.00 | 0.00 | 0.00 | 0.00 | 0.00 | 0.00 | 0.00 | 0.00 | 0.00 | 0.00 | 0.00 | 0.00 | 0.00 |
| | Patient22 | 1.83 | 0.00 | 0.00 | 0.00 | 0.00 | 0.00 | 0.00 | 0.00 | 0.00 | 0.00 | 0.00 | 0.00 | 0.00 | 0.00 | 0.00 | 0.00 |
| | Patient23 | 1.28 | 0.00 | 0.28 | 0.00 | 0.00 | 0.00 | 0.42 | 0.00 | 0.59 | 0.00 | 0.00 | 0.00 | 0.00 | 0.00 | 0.00 | 0.00 |
| | Patient24 | 1.38 | 0.00 | 0.00 | 0.00 | 0.00 | 0.00 | 0.00 | 0.00 | 0.39 | 0.00 | 0.00 | 0.11 | 0.00 | 0.00 | 0.00 | 0.00 |
| | Patient25 | 0.56 | 0.00 | 0.00 | 0.00 | 0.03 | 0.00 | 0.00 | 0.00 | 0.00 | 0.00 | 0.00 | 0.00 | 0.00 | 0.00 | 0.00 | 0.00 |
| | Patient26 | 0.91 | 0.00 | 0.21 | 0.66 | 0.00 | 0.00 | 0.23 | 0.00 | 0.23 | 0.00 | 0.16 | 0.00 | 0.00 | 0.00 | 0.00 | 0.66 |
| | Patient27 | 0.99 | 0.00 | 0.00 | 0.00 | 0.00 | 0.00 | 0.00 | 0.00 | 0.00 | 0.00 | 0.00 | 0.00 | 0.16 | 0.00 | 0.00 | 0.00 |
| | Patient28 | 0.69 | 0.00 | 0.00 | 0.00 | 0.00 | 0.00 | 0.00 | 0.00 | 0.00 | 0.00 | 0.00 | 0.00 | 0.00 | 0.00 | 0.00 | 0.58 |
| | Patient29 | 1.41 | 0.00 | 0.00 | 0.00 | 0.00 | 0.00 | 0.62 | 0.00 | 0.63 | 0.00 | 0.00 | 0.00 | 0.00 | 0.00 | 0.00 | 0.00 |
| | Patient30 | 1.55 | 0.00 | 0.01 | 0.00 | 0.00 | 0.00 | 0.00 | 0.00 | 0.52 | 0.00 | 0.00 | 0.00 | 0.00 | 0.00 | 0.00 | 0.00 |

| | | Nonsense mediated decay | | | | | | | | | | | | | | | | | | |
|---|---|---|---|---|---|---|---|---|---|---|---|---|---|---|---|---|---|---|---|---|
| | | ATXN3-202 | ATXN3-212 | ATXN3-218 | ATXN3-221 | ATXN3-225 | ATXN3-229 | ATXN3-230 | ATXN3-232 | ATXN3-234 | ATXN3-236 | ATXN3-237 | ATXN3-238 | ATXN3-240 | ATXN3-242 | ATXN3-243 | ATXN3-244 | ATXN3-245 | ATXN3-253 | ATXN3-254 |
| Cerebellum | CTRL1 | 0.08 | 0.00 | 0.00 | 0.00 | 0.38 | 0.00 | 0.00 | 0.00 | 3.77 | 0.00 | 0.00 | 0.00 | 0.00 | 0.00 | 0.95 | 0.00 | 0.00 | 0.63 | 0.14 |
| | CTRL2 | 0.01 | 0.00 | 0.00 | 0.00 | 0.21 | 0.00 | 0.37 | 0.00 | 3.27 | 0.00 | 0.36 | 0.00 | 0.00 | 0.00 | 0.80 | 1.02 | 0.00 | 0.00 | 0.06 |
| | CTRL3 | 0.03 | 0.00 | 0.00 | 0.00 | 0.16 | 0.00 | 2.35 | 0.00 | 5.16 | 0.00 | 0.14 | 0.00 | 0.36 | 0.00 | 0.00 | 0.57 | 0.00 | 0.32 | 0.17 |
| | CTRL4 | 0.00 | 0.00 | 0.00 | 0.00 | 0.00 | 0.00 | 0.00 | 0.00 | 1.75 | 0.00 | 0.25 | 0.91 | 0.69 | 0.00 | 4.50 | 0.00 | 0.00 | 0.00 | 0.00 |
| | CTRL5 | 0.00 | 0.00 | 0.00 | 0.00 | 0.00 | 0.00 | 0.00 | 1.27 | 0.51 | 0.00 | 0.00 | 0.00 | 0.00 | 0.00 | 0.93 | 0.91 | 0.42 | 0.15 | 0.00 |
| | CTRL6 | 0.00 | 0.00 | 0.00 | 0.00 | 0.00 | 0.10 | 0.35 | 0.00 | 4.24 | 0.00 | 0.00 | 0.00 | 0.07 | 0.00 | 0.00 | 0.00 | 0.00 | 0.24 | 0.07 |
| | Patient1 | 0.00 | 0.00 | 0.00 | 0.00 | 0.00 | 0.00 | 0.00 | 0.00 | 2.75 | 0.00 | 0.00 | 0.00 | 0.00 | 0.00 | 1.04 | 0.00 | 0.00 | 0.31 | 0.04 |
| | Patient2 | 0.00 | 0.00 | 0.00 | 0.00 | 0.00 | 0.00 | 0.00 | 1.75 | 0.00 | 0.00 | 0.86 | 0.65 | 0.00 | 0.00 | 1.99 | 1.26 | 0.00 | 0.48 | 0.06 |
| | Patient3 | 0.00 | 0.00 | 0.00 | 0.00 | 0.00 | 0.00 | 0.00 | 0.00 | 2.50 | 0.00 | 0.00 | 0.00 | 0.00 | 1.92 | 0.22 | 0.00 | 0.00 | 0.00 | 0.13 |
| | Patient4 | 0.00 | 0.00 | 0.00 | 0.00 | 0.38 | 0.00 | 0.00 | 0.00 | 6.06 | 0.00 | 0.00 | 0.00 | 0.00 | 0.00 | 1.67 | 0.00 | 0.30 | 0.00 | 0.00 |
| | Patient5 | 0.00 | 0.00 | 0.00 | 0.00 | 0.44 | 0.00 | 0.00 | 0.00 | 7.36 | 0.00 | 0.00 | 0.00 | 0.29 | 0.00 | 0.00 | 0.00 | 0.00 | 0.14 | 0.20 |
| | Patient6 | 0.03 | 0.00 | 0.00 | 0.00 | 0.41 | 0.00 | 0.20 | 0.62 | 2.28 | 0.00 | 0.00 | 0.11 | 0.03 | 0.00 | 1.65 | 0.00 | 0.34 | 0.26 | 0.00 |
| Blood | CTRL1 | 0.20 | 0.00 | 0.13 | 0.00 | 0.00 | 0.00 | 0.00 | 0.00 | 0.01 | 0.00 | 0.00 | 0.00 | 0.00 | 0.00 | 0.00 | 0.00 | 0.00 | 0.14 | 0.11 |
| | CTRL2 | 0.53 | 0.00 | 0.00 | 0.00 | 0.00 | 0.00 | 0.00 | 0.00 | 0.00 | 0.00 | 0.13 | 0.00 | 0.00 | 0.00 | 0.00 | 0.00 | 0.00 | 0.00 | 0.00 |
| | CTRL3 | 0.38 | 0.00 | 0.00 | 0.00 | 0.00 | 0.00 | 0.00 | 0.00 | 0.00 | 0.00 | 0.00 | 0.00 | 0.00 | 0.00 | 0.00 | 0.00 | 0.00 | 0.00 | 0.06 |
| | CTRL4 | 0.25 | 0.00 | 0.00 | 0.00 | 0.00 | 0.00 | 0.00 | 0.00 | 0.00 | 0.00 | 0.00 | 0.00 | 0.00 | 0.00 | 0.00 | 0.00 | 0.00 | 0.00 | 0.03 |
| | CTRL5 | 0.04 | 0.00 | 0.00 | 0.00 | 0.00 | 0.00 | 0.00 | 0.00 | 0.00 | 0.00 | 0.00 | 0.00 | 0.00 | 0.00 | 0.00 | 0.00 | 0.00 | 0.00 | 0.00 |
| | CTRL6 | 0.53 | 0.00 | 0.00 | 0.00 | 0.00 | 0.09 | 0.00 | 0.00 | 0.12 | 0.00 | 0.00 | 0.00 | 0.00 | 0.00 | 0.00 | 0.00 | 0.00 | 0.00 | 0.00 |
| | CTRL7 | 0.66 | 0.00 | 0.00 | 0.00 | 0.00 | 0.00 | 0.00 | 0.00 | 0.26 | 0.00 | 0.00 | 0.00 | 0.00 | 0.00 | 0.00 | 0.00 | 0.00 | 0.00 | 0.19 |
| | CTRL8 | 0.29 | 0.00 | 0.00 | 0.00 | 0.00 | 0.00 | 0.08 | 0.00 | 0.00 | 0.00 | 0.00 | 0.00 | 0.00 | 0.00 | 0.00 | 0.00 | 0.00 | 0.00 | 0.00 |
| | CTRL9 | 1.79 | 0.00 | 0.00 | 0.00 | 0.00 | 0.12 | 0.00 | 0.00 | 0.00 | 0.00 | 0.20 | 0.00 | 0.00 | 0.00 | 0.00 | 0.00 | 0.00 | 0.19 | 0.20 |
| | CTRL10 | 1.27 | 0.00 | 0.00 | 0.00 | 0.00 | 0.28 | 0.00 | 0.00 | 0.00 | 0.00 | 0.00 | 0.00 | 0.00 | 0.00 | 0.00 | 0.00 | 0.00 | 0.00 | 0.00 |
| | CTRL11 | 0.36 | 0.00 | 0.00 | 0.00 | 0.00 | 0.00 | 0.00 | 0.00 | 0.00 | 0.00 | 0.00 | 0.00 | 0.00 | 0.00 | 0.00 | 0.00 | 0.00 | 0.37 | 0.00 |
| | CTRL12 | 1.13 | 0.06 | 0.00 | 0.00 | 0.00 | 0.24 | 0.00 | 0.00 | 0.00 | 0.00 | 0.00 | 0.60 | 0.00 | 0.00 | 0.00 | 0.00 | 0.00 | 0.00 | 0.06 |
| | CTRL13 | 1.59 | 0.00 | 0.00 | 0.17 | 0.00 | 0.00 | 0.00 | 0.00 | 0.00 | 0.00 | 0.00 | 0.00 | 0.00 | 0.00 | 0.00 | 0.00 | 0.00 | 0.00 | 0.00 |
| | CTRL14 | 0.47 | 0.00 | 0.00 | 0.00 | 0.00 | 0.00 | 0.00 | 0.00 | 0.04 | 0.00 | 0.00 | 0.00 | 0.00 | 0.00 | 0.00 | 0.00 | 0.00 | 0.00 | 0.00 |
| | CTRL15 | 1.31 | 0.00 | 0.00 | 0.00 | 0.00 | 0.13 | 0.00 | 0.00 | 0.00 | 0.00 | 0.00 | 0.00 | 0.00 | 0.00 | 0.00 | 0.00 | 0.00 | 0.23 | 0.00 |
| | CTRL16 | 0.80 | 0.00 | 0.00 | 0.00 | 0.00 | 0.00 | 0.00 | 0.00 | 0.00 | 0.00 | 0.11 | 0.00 | 0.00 | 0.00 | 0.12 | 0.00 | 0.00 | 0.00 | 0.21 |
| | CTRL17 | 1.22 | 0.00 | 0.00 | 0.00 | 0.00 | 0.19 | 0.00 | 0.00 | 0.00 | 0.00 | 0.19 | 0.00 | 0.00 | 0.00 | 0.43 | 0.00 | 0.00 | 0.00 | 0.00 |
| | CTRL18 | 2.05 | 0.00 | 0.00 | 0.00 | 0.00 | 0.00 | 0.00 | 0.00 | 0.00 | 0.00 | 0.00 | 0.00 | 0.00 | 0.00 | 0.00 | 0.00 | 0.00 | 0.00 | 0.00 |
| | CTRL19 | 0.55 | 0.00 | 0.00 | 0.00 | 0.00 | 0.30 | 0.00 | 0.00 | 0.00 | 0.00 | 0.00 | 0.00 | 0.00 | 0.00 | 0.00 | 0.00 | 0.00 | 0.00 | 0.09 |
| | CTRL20 | 0.39 | 0.00 | 0.00 | 0.00 | 0.00 | 0.00 | 0.00 | 0.00 | 0.07 | 0.00 | 0.00 | 0.00 | 0.00 | 0.00 | 0.00 | 0.00 | 0.00 | 0.00 | 0.00 |
| | Patient1 | 0.59 | 0.00 | 0.00 | 0.00 | 0.18 | 0.00 | 0.00 | 0.00 | 0.00 | 0.00 | 0.00 | 0.00 | 0.00 | 0.00 | 0.00 | 0.00 | 0.00 | 0.00 | 0.19 |
| | Patient2 | 0.18 | 0.00 | 0.00 | 0.00 | 0.00 | 0.00 | 0.00 | 0.00 | 0.00 | 0.00 | 0.00 | 0.00 | 0.00 | 0.00 | 0.00 | 0.00 | 0.00 | 0.00 | 0.11 |
| | Patient3 | 0.57 | 0.00 | 0.00 | 0.00 | 0.00 | 0.13 | 0.00 | 0.00 | 0.00 | 0.00 | 0.00 | 0.00 | 0.00 | 0.00 | 0.00 | 0.00 | 0.00 | 0.00 | 0.00 |
| | Patient4 | 0.45 | 0.00 | 0.00 | 0.00 | 0.00 | 0.00 | 0.00 | 0.00 | 0.00 | 0.00 | 0.26 | 0.00 | 0.00 | 0.00 | 0.00 | 0.00 | 0.00 | 0.00 | 0.16 |
| | Patient5 | 0.41 | 0.00 | 0.00 | 0.00 | 0.00 | 0.00 | 0.00 | 0.00 | 0.00 | 0.00 | 0.00 | 0.00 | 0.00 | 0.00 | 0.00 | 0.00 | 0.00 | 0.24 | 0.12 |
| | Patient6 | 0.49 | 0.00 | 0.00 | 0.00 | 0.00 | 0.00 | 0.00 | 0.00 | 0.00 | 0.00 | 0.00 | 0.00 | 0.00 | 0.00 | 0.00 | 0.00 | 0.00 | 0.00 | 0.00 |
| | Patient7 | 0.37 | 0.00 | 0.00 | 0.00 | 0.00 | 0.28 | 0.00 | 0.00 | 0.05 | 0.00 | 0.00 | 0.00 | 0.00 | 0.00 | 0.00 | 0.00 | 0.00 | 0.00 | 0.19 |
| | Patient8 | 1.14 | 0.00 | 0.00 | 0.00 | 0.00 | 0.58 | 0.00 | 0.05 | 0.00 | 0.00 | 0.00 | 0.00 | 0.00 | 0.00 | 0.00 | 0.00 | 0.00 | 0.00 | 0.00 |
| | Patient9 | 1.81 | 0.00 | 0.00 | 0.00 | 0.00 | 0.00 | 0.00 | 0.00 | 0.00 | 0.00 | 0.00 | 0.00 | 0.00 | 0.00 | 0.00 | 0.00 | 0.00 | 0.00 | 0.10 |
| | Patient10 | 0.16 | 0.00 | 0.00 | 0.00 | 0.00 | 0.00 | 0.00 | 0.00 | 0.00 | 0.00 | 0.00 | 0.00 | 0.00 | 0.00 | 0.00 | 0.00 | 0.00 | 0.00 | 0.06 |
| | Patient11 | 0.00 | 0.00 | 0.00 | 0.00 | 0.00 | 0.00 | 0.00 | 0.00 | 0.00 | 0.00 | 0.00 | 0.00 | 0.00 | 0.00 | 0.00 | 0.00 | 0.00 | 0.00 | 0.10 |
| | Patient12 | 0.13 | 0.00 | 0.00 | 0.00 | 0.00 | 0.00 | 0.00 | 0.00 | 0.00 | 0.00 | 0.16 | 0.00 | 0.00 | 0.00 | 0.15 | 0.00 | 0.00 | 0.00 | 0.00 |
| | Patient13 | 0.15 | 0.00 | 0.00 | 0.00 | 0.00 | 0.00 | 0.00 | 0.00 | 0.00 | 0.00 | 0.00 | 0.00 | 0.00 | 0.00 | 0.00 | 0.00 | 0.00 | 0.00 | 0.00 |
| | Patient14 | 0.26 | 0.00 | 0.00 | 0.00 | 0.00 | 0.00 | 0.00 | 0.00 | 0.00 | 0.00 | 0.00 | 0.00 | 0.00 | 0.00 | 0.00 | 0.00 | 0.00 | 0.00 | 0.07 |
| | Patient15 | 0.66 | 0.00 | 0.00 | 0.00 | 0.00 | 0.00 | 0.00 | 0.00 | 0.00 | 0.00 | 0.00 | 0.00 | 0.00 | 0.00 | 0.00 | 0.00 | 0.00 | 0.00 | 0.00 |
| | Patient16 | 1.05 | 0.00 | 0.00 | 0.00 | 0.00 | 0.00 | 0.00 | 0.00 | 0.00 | 0.00 | 0.00 | 0.00 | 0.00 | 0.00 | 0.25 | 0.00 | 0.00 | 0.00 | 0.18 |
| | Patient17 | 0.54 | 0.00 | 0.00 | 0.00 | 0.00 | 0.00 | 0.00 | 0.02 | 0.00 | 0.00 | 0.00 | 0.00 | 0.00 | 0.00 | 0.08 | 0.00 | 0.00 | 0.00 | 0.08 |
| | Patient18 | 1.21 | 0.00 | 0.00 | 0.00 | 0.00 | 0.00 | 0.00 | 0.00 | 0.00 | 0.00 | 0.00 | 0.00 | 0.00 | 0.00 | 0.00 | 0.00 | 0.00 | 0.00 | 0.00 |
| | Patient19 | 0.82 | 0.00 | 0.00 | 0.00 | 0.00 | 0.00 | 0.00 | 0.00 | 0.00 | 0.00 | 0.00 | 0.00 | 0.00 | 0.00 | 0.00 | 0.00 | 0.00 | 0.00 | 0.00 |
| | Patient20 | 1.55 | 0.00 | 0.00 | 0.00 | 0.00 | 0.11 | 0.00 | 0.00 | 0.00 | 0.00 | 0.00 | 0.00 | 0.00 | 0.00 | 0.00 | 0.00 | 0.00 | 0.39 | 0.12 |
| | Patient21 | 0.44 | 0.00 | 0.00 | 0.00 | 0.00 | 0.00 | 0.00 | 0.00 | 0.00 | 0.00 | 0.07 | 0.00 | 0.00 | 0.00 | 0.00 | 0.00 | 0.00 | 0.00 | 0.00 |
| | Patient22 | 1.44 | 0.00 | 0.00 | 0.00 | 0.00 | 0.26 | 0.00 | 0.00 | 0.00 | 0.00 | 0.00 | 0.35 | 0.00 | 0.00 | 0.00 | 0.00 | 0.00 | 0.16 | 0.16 |
| | Patient23 | 1.42 | 0.00 | 0.00 | 0.24 | 0.00 | 0.00 | 0.00 | 0.00 | 0.00 | 0.00 | 0.00 | 0.00 | 0.00 | 0.00 | 0.00 | 0.00 | 0.00 | 0.00 | 0.00 |
| | Patient24 | 1.14 | 0.00 | 0.00 | 0.00 | 0.00 | 0.00 | 0.00 | 0.00 | 0.00 | 0.00 | 0.00 | 0.00 | 0.00 | 0.00 | 0.00 | 0.00 | 0.00 | 0.00 | 0.07 |
| | Patient25 | 0.29 | 0.00 | 0.00 | 0.00 | 0.00 | 0.00 | 0.00 | 0.00 | 0.00 | 0.00 | 0.00 | 0.32 | 0.00 | 0.00 | 0.00 | 0.00 | 0.00 | 0.00 | 0.22 |
| | Patient26 | 1.12 | 0.00 | 0.00 | 0.00 | 0.00 | 0.34 | 0.00 | 0.00 | 0.00 | 0.00 | 0.00 | 0.00 | 0.00 | 0.00 | 0.00 | 0.00 | 0.00 | 0.31 | 0.00 |
| | Patient27 | 0.74 | 0.00 | 0.00 | 0.00 | 0.00 | 0.00 | 0.00 | 0.00 | 0.00 | 0.14 | 0.00 | 0.00 | 0.00 | 0.00 | 0.22 | 0.00 | 0.00 | 0.00 | 0.13 |
| | Patient28 | 0.32 | 0.00 | 0.00 | 0.00 | 0.00 | 0.00 | 0.00 | 0.00 | 0.00 | 0.00 | 0.00 | 0.00 | 0.00 | 0.00 | 0.00 | 0.00 | 0.00 | 0.00 | 0.45 |
| | Patient29 | 1.97 | 0.00 | 0.00 | 0.00 | 0.00 | 0.00 | 0.00 | 0.00 | 0.00 | 0.00 | 0.00 | 0.00 | 0.00 | 0.00 | 0.00 | 0.00 | 0.00 | 0.00 | 0.19 |
| | Patient30 | 1.19 | 0.27 | 0.00 | 0.00 | 0.00 | 0.00 | 0.00 | 0.00 | 0.64 | 0.00 | 0.00 | 0.00 | 0.00 | 0.00 | 0.00 | 0.00 | 0.00 | 0.00 | 0.21 |

| | | Protein CDS not defined | | | | | | | | | | | | | | | | Retained intron | | | * | | |
|---|---|---|---|---|---|---|---|---|---|---|---|---|---|---|---|---|---|---|---|---|---|---|---|
| | | ATXN3-208 | ATXN3-211 | ATXN3-216 | ATXN3-217 | ATXN3-220 | ATXN3-222 | ATXN3-223 | ATXN3-224 | ATXN3-226 | ATXN3-233 | ATXN3-239 | ATXN3-241 | ATXN3-247 | ATXN3-249 | ATXN3-250 | ATXN3-252 | ATXN3-205 | ATXN3-210 | ATXN3-248 | ENST00000617719 | ENST00000620536 | ENST00000621269 |
| Cerebellum | CTRL1 | 20.94 | 0.00 | 0.00 | 1.34 | 0.00 | 0.00 | 0.00 | 0.00 | 0.29 | 0.00 | 4.44 | 0.00 | 0.00 | 0.00 | 0.00 | 0.00 | 0.23 | 0.00 | 0.00 | 0.00 | 0.00 | 0.00 |
| | CTRL2 | 26.05 | 0.32 | 0.00 | 2.12 | 0.00 | 0.00 | 0.00 | 0.00 | 0.67 | 0.00 | 0.00 | 0.37 | 0.00 | 0.00 | 0.00 | 0.00 | 0.80 | 0.10 | 0.03 | 0.38 | 0.00 | 0.42 |
| | CTRL3 | 38.25 | 0.00 | 0.31 | 0.00 | 0.00 | 0.00 | 0.00 | 0.18 | 0.70 | 0.00 | 0.00 | 0.00 | 0.00 | 0.00 | 0.00 | 0.00 | 0.72 | 0.29 | 0.00 | 0.00 | 0.00 | 5.01 |
| | CTRL4 | 50.49 | 0.17 | 0.00 | 1.55 | 0.00 | 0.00 | 0.00 | 0.00 | 0.04 | 0.00 | 0.00 | 0.00 | 0.00 | 0.00 | 0.00 | 0.00 | 3.80 | 0.00 | 0.00 | 0.64 | 0.00 | 0.00 |
| | CTRL5 | 29.52 | 0.00 | 0.00 | 1.07 | 0.00 | 0.00 | 0.00 | 0.00 | 0.90 | 0.00 | 0.00 | 0.00 | 0.00 | 0.00 | 0.00 | 0.00 | 0.31 | 0.15 | 0.00 | 0.00 | 0.00 | 2.82 |
| | CTRL6 | 35.41 | 0.00 | 0.00 | 1.49 | 0.00 | 0.00 | 0.00 | 0.00 | 0.82 | 0.00 | 0.00 | 0.00 | 0.00 | 0.00 | 0.05 | 0.00 | 0.93 | 0.15 | 0.00 | 0.17 | 0.00 | 1.26 |
| | Patient1 | 44.92 | 0.00 | 0.00 | 0.67 | 0.00 | 0.00 | 0.00 | 0.00 | 1.58 | 0.00 | 0.00 | 0.00 | 0.00 | 0.02 | 0.00 | 0.00 | 0.66 | 0.00 | 0.04 | 0.00 | 0.00 | 2.35 |
| | Patient2 | 39.26 | 0.00 | 0.00 | 0.00 | 0.00 | 0.00 | 0.00 | 0.00 | 0.93 | 0.00 | 0.00 | 0.00 | 0.00 | 0.00 | 0.40 | 0.00 | 0.09 | 0.42 | 0.00 | 0.00 | 0.00 | 0.00 |
| | Patient3 | 50.20 | 0.00 | 0.00 | 3.23 | 0.00 | 4.21 | 0.00 | 0.00 | 0.00 | 0.00 | 0.00 | 0.00 | 0.00 | 0.00 | 0.00 | 0.17 | 0.55 | 0.21 | 0.00 | 0.00 | 0.00 | 0.00 |
| | Patient4 | 38.87 | 0.00 | 0.00 | 0.34 | 0.00 | 0.00 | 0.16 | 0.00 | 0.00 | 0.00 | 0.00 | 0.00 | 0.00 | 0.00 | 0.00 | 0.00 | 1.05 | 0.25 | 0.00 | 0.53 | 0.00 | 1.36 |
| | Patient5 | 24.65 | 0.00 | 0.00 | 0.95 | 0.00 | 0.00 | 0.00 | 0.77 | 0.00 | 0.00 | 0.00 | 0.00 | 0.00 | 0.00 | 0.00 | 0.00 | 0.35 | 0.11 | 0.00 | 0.00 | 0.00 | 0.00 |
| | Patient6 | 40.78 | 0.00 | 0.00 | 2.09 | 0.00 | 0.00 | 0.45 | 0.00 | 0.00 | 0.00 | 0.00 | 0.00 | 0.08 | 0.00 | 0.00 | 0.00 | 0.50 | 0.42 | 0.00 | 0.00 | 0.00 | 2.23 |
| Blood | CTRL1 | 0.00 | 0.00 | 0.00 | 0.00 | 0.04 | 0.00 | 0.00 | 0.00 | 0.00 | 0.00 | 0.00 | 0.00 | 0.00 | 0.00 | 0.00 | 0.00 | 0.00 | 0.27 | 0.26 | 0.00 | 0.00 | 0.00 |
| | CTRL2 | 0.00 | 0.00 | 0.00 | 0.00 | 0.00 | 0.00 | 0.00 | 0.00 | 0.00 | 0.00 | 0.00 | 0.00 | 0.00 | 0.08 | 0.25 | 0.00 | 0.09 | 0.25 | 0.05 | 0.22 | 0.00 | 0.00 |
| | CTRL3 | 0.00 | 0.00 | 0.00 | 0.00 | 0.00 | 0.00 | 0.00 | 0.09 | 0.00 | 0.00 | 0.00 | 0.00 | 0.00 | 0.00 | 0.00 | 0.00 | 0.25 | 0.13 | 0.11 | 0.00 | 0.00 | 0.00 |
| | CTRL4 | 0.00 | 0.00 | 0.00 | 0.00 | 0.00 | 0.00 | 0.00 | 0.00 | 0.06 | 0.00 | 0.00 | 0.00 | 0.00 | 0.03 | 0.00 | 0.00 | 0.00 | 0.07 | 0.00 | 0.00 | 0.00 | 0.00 |
| | CTRL5 | 0.00 | 0.00 | 0.00 | 0.00 | 0.00 | 0.00 | 0.00 | 0.00 | 0.00 | 0.00 | 0.00 | 0.34 | 0.00 | 0.00 | 0.00 | 0.00 | 0.00 | 0.00 | 0.00 | 0.00 | 0.00 | 0.00 |
| | CTRL6 | 0.00 | 0.25 | 0.00 | 0.00 | 0.00 | 0.00 | 0.00 | 0.00 | 0.00 | 0.17 | 0.00 | 0.00 | 0.31 | 0.00 | 0.00 | 0.00 | 0.07 | 0.90 | 0.14 | 0.00 | 0.00 | 0.05 |
| | CTRL7 | 0.00 | 0.10 | 0.00 | 0.00 | 0.00 | 0.00 | 0.00 | 0.62 | 0.00 | 0.00 | 0.00 | 0.00 | 0.00 | 0.04 | 0.00 | 0.00 | 0.07 | 0.48 | 0.22 | 0.00 | 0.00 | 0.04 |
| | CTRL8 | 0.00 | 0.14 | 0.00 | 0.00 | 0.08 | 0.00 | 0.00 | 0.00 | 0.12 | 0.00 | 0.00 | 0.00 | 0.00 | 0.05 | 0.53 | 0.00 | 0.00 | 0.08 | 0.00 | 0.00 | 0.00 | 0.00 |
| | CTRL9 | 0.00 | 0.00 | 0.00 | 0.00 | 0.00 | 0.00 | 0.00 | 0.00 | 0.00 | 0.00 | 0.00 | 0.00 | 0.00 | 0.28 | 0.00 | 0.00 | 0.16 | 0.47 | 0.23 | 0.00 | 0.00 | 0.00 |
| | CTRL10 | 0.00 | 0.00 | 0.00 | 0.00 | 0.00 | 0.00 | 0.00 | 0.00 | 0.00 | 0.00 | 0.00 | 0.00 | 0.00 | 0.41 | 0.12 | 0.00 | 0.34 | 0.17 | 0.06 | 0.00 | 0.00 | 1.96 |
| | CTRL11 | 0.27 | 0.00 | 0.00 | 0.00 | 0.00 | 0.00 | 0.00 | 0.00 | 0.33 | 0.00 | 0.00 | 0.00 | 0.00 | 0.04 | 0.00 | 0.00 | 0.00 | 0.30 | 0.16 | 0.00 | 0.00 | 0.00 |
| | CTRL12 | 0.00 | 0.00 | 0.00 | 0.00 | 0.00 | 0.00 | 0.00 | 0.00 | 0.00 | 0.00 | 0.00 | 0.00 | 0.00 | 0.11 | 0.20 | 0.00 | 0.09 | 1.18 | 0.33 | 0.00 | 0.00 | 0.58 |
| | CTRL13 | 0.00 | 0.00 | 0.00 | 0.00 | 0.00 | 0.00 | 0.00 | 0.00 | 0.00 | 0.00 | 0.00 | 0.00 | 0.00 | 0.09 | 0.35 | 0.00 | 0.00 | 0.72 | 0.35 | 0.00 | 0.00 | 0.80 |
| | CTRL14 | 0.31 | 0.00 | 0.00 | 0.00 | 0.00 | 0.00 | 0.00 | 0.00 | 0.00 | 0.00 | 0.00 | 0.00 | 0.09 | 0.11 | 0.00 | 0.00 | 0.10 | 0.26 | 0.29 | 0.00 | 0.00 | 0.00 |
| | CTRL15 | 0.00 | 0.00 | 0.21 | 0.00 | 0.23 | 0.00 | 0.00 | 0.00 | 0.02 | 0.00 | 0.00 | 0.00 | 0.00 | 0.00 | 0.12 | 0.00 | 0.00 | 0.21 | 0.13 | 0.00 | 0.00 | 0.74 |
| | CTRL16 | 0.00 | 0.00 | 0.00 | 0.00 | 0.00 | 0.00 | 0.00 | 0.00 | 0.00 | 0.00 | 0.00 | 0.00 | 0.00 | 0.06 | 0.00 | 0.00 | 0.07 | 0.29 | 0.06 | 0.15 | 0.00 | 0.00 |
| | CTRL17 | 0.00 | 0.26 | 0.00 | 0.00 | 0.00 | 0.00 | 0.00 | 0.32 | 0.34 | 0.00 | 0.00 | 0.00 | 0.00 | 0.07 | 0.00 | 0.21 | 0.09 | 0.57 | 0.49 | 0.00 | 0.00 | 0.48 |
| | CTRL18 | 0.47 | 0.16 | 0.00 | 0.00 | 0.27 | 0.00 | 0.00 | 0.21 | 0.23 | 0.00 | 0.00 | 0.00 | 0.49 | 0.00 | 0.00 | 0.00 | 0.23 | 0.46 | 0.31 | 0.00 | 0.00 | 0.00 |
| | CTRL19 | 0.00 | 0.00 | 0.00 | 0.00 | 0.00 | 0.00 | 0.00 | 0.00 | 0.00 | 0.18 | 0.02 | 0.00 | 0.00 | 0.00 | 0.00 | 0.00 | 0.00 | 0.00 | 0.16 | 0.00 | 0.00 | 0.19 |
| | CTRL20 | 0.00 | 0.00 | 0.00 | 0.00 | 0.00 | 0.00 | 0.00 | 0.00 | 0.00 | 0.42 | 0.00 | 0.00 | 0.00 | 0.00 | 0.12 | 0.00 | 0.13 | 0.00 | 0.00 | 0.22 | 0.00 | 0.00 |
| | Patient1 | 0.00 | 0.00 | 0.00 | 0.00 | 0.00 | 0.00 | 0.00 | 0.00 | 0.00 | 0.02 | 0.00 | 0.00 | 0.14 | 0.00 | 0.00 | 0.00 | 0.00 | 0.09 | 0.00 | 0.00 | 0.00 | 0.38 |
| | Patient2 | 0.00 | 0.00 | 0.00 | 0.00 | 0.00 | 0.00 | 0.00 | 0.00 | 0.00 | 0.00 | 0.00 | 0.00 | 0.00 | 0.00 | 0.00 | 0.00 | 0.00 | 0.07 | 0.00 | 0.00 | 0.00 | 0.00 |
| | Patient3 | 0.00 | 0.00 | 0.02 | 0.04 | 0.00 | 0.00 | 0.00 | 0.13 | 0.00 | 0.00 | 0.00 | 0.00 | 0.17 | 0.00 | 0.00 | 0.00 | 0.08 | 0.30 | 0.16 | 0.00 | 0.00 | 0.00 |
| | Patient4 | 0.20 | 0.00 | 0.00 | 0.00 | 0.00 | 0.00 | 0.00 | 0.00 | 0.00 | 0.00 | 0.00 | 0.00 | 0.00 | 0.04 | 0.00 | 0.00 | 0.00 | 0.48 | 0.33 | 0.00 | 0.00 | 0.00 |
| | Patient5 | 0.00 | 0.00 | 0.00 | 0.00 | 0.00 | 0.00 | 0.00 | 0.00 | 0.13 | 0.00 | 0.00 | 0.00 | 0.00 | 0.12 | 0.00 | 0.00 | 0.09 | 0.34 | 0.00 | 0.00 | 0.00 | 0.00 |
| | Patient6 | 0.00 | 0.00 | 0.27 | 0.00 | 0.00 | 0.00 | 0.00 | 0.00 | 0.00 | 0.12 | 0.00 | 0.00 | 0.00 | 0.08 | 0.00 | 0.00 | 0.00 | 0.00 | 0.00 | 0.00 | 0.00 | 0.00 |
| | Patient7 | 0.60 | 0.00 | 0.00 | 0.00 | 0.00 | 0.00 | 0.00 | 0.00 | 0.00 | 0.00 | 0.00 | 0.00 | 0.00 | 0.00 | 0.00 | 0.00 | 0.11 | 0.13 | 0.24 | 0.00 | 0.00 | 0.68 |
| | Patient8 | 0.21 | 0.19 | 0.00 | 0.00 | 0.00 | 0.00 | 0.00 | 0.00 | 0.00 | 0.00 | 0.00 | 0.00 | 0.40 | 0.16 | 0.00 | 0.00 | 0.36 | 0.32 | 0.00 | 0.00 | 0.00 | 0.52 |
| | Patient9 | 0.00 | 0.43 | 0.00 | 0.04 | 0.00 | 0.62 | 0.20 | 0.15 | 0.00 | 0.00 | 0.00 | 0.00 | 0.00 | 0.10 | 0.00 | 0.00 | 0.00 | 0.32 | 0.00 | 0.00 | 0.00 | 1.45 |
| | Patient10 | 0.00 | 0.00 | 0.00 | 0.00 | 0.00 | 0.00 | 0.00 | 0.00 | 0.00 | 0.00 | 0.00 | 0.00 | 0.00 | 0.03 | 0.00 | 0.00 | 0.00 | 0.00 | 0.00 | 0.00 | 0.00 | 0.18 |
| | Patient11 | 0.00 | 0.00 | 0.00 | 0.00 | 0.10 | 0.00 | 0.00 | 0.00 | 0.00 | 0.00 | 0.00 | 0.00 | 0.00 | 0.08 | 0.17 | 0.00 | 0.08 | 0.45 | 0.13 | 0.00 | 0.00 | 0.00 |
| | Patient12 | 0.00 | 0.12 | 0.00 | 0.00 | 0.00 | 0.00 | 0.00 | 0.00 | 0.20 | 0.00 | 0.00 | 0.00 | 0.44 | 0.05 | 0.00 | 0.00 | 0.00 | 0.69 | 0.00 | 0.00 | 0.00 | 0.00 |
| | Patient13 | 0.00 | 0.00 | 0.00 | 0.00 | 0.00 | 0.00 | 0.12 | 0.00 | 0.00 | 0.00 | 0.00 | 0.27 | 0.00 | 0.04 | 0.00 | 0.00 | 0.09 | 0.29 | 0.11 | 0.00 | 0.00 | 0.00 |
| | Patient14 | 0.00 | 0.00 | 0.00 | 0.00 | 0.00 | 0.00 | 0.00 | 0.00 | 0.00 | 0.00 | 0.00 | 0.00 | 0.00 | 0.07 | 0.04 | 0.00 | 0.00 | 0.00 | 0.00 | 0.00 | 0.00 | 0.00 |
| | Patient15 | 0.49 | 0.00 | 0.00 | 0.00 | 0.00 | 0.00 | 0.00 | 0.00 | 0.00 | 0.00 | 0.00 | 0.00 | 0.00 | 0.00 | 0.00 | 0.00 | 0.00 | 0.44 | 0.12 | 0.00 | 0.00 | 0.16 |
| | Patient16 | 0.00 | 0.00 | 0.00 | 0.00 | 0.00 | 0.35 | 0.00 | 0.00 | 0.00 | 0.59 | 0.00 | 0.00 | 0.00 | 0.07 | 0.00 | 0.09 | 0.00 | 0.60 | 0.00 | 0.00 | 0.00 | 0.00 |
| | Patient17 | 0.00 | 0.00 | 0.28 | 0.00 | 0.00 | 0.00 | 0.00 | 0.00 | 0.00 | 0.00 | 0.00 | 0.00 | 0.11 | 0.00 | 0.00 | 0.00 | 0.11 | 0.41 | 0.06 | 0.00 | 0.00 | 0.00 |
| | Patient18 | 1.25 | 0.27 | 0.00 | 0.00 | 0.00 | 0.00 | 0.00 | 0.34 | 0.00 | 0.17 | 0.00 | 0.00 | 0.00 | 0.32 | 0.00 | 0.08 | 0.10 | 0.00 | 0.23 | 0.00 | 0.00 | 0.19 |
| | Patient19 | 0.61 | 0.23 | 0.00 | 0.52 | 0.00 | 0.00 | 0.00 | 0.00 | 0.00 | 0.00 | 0.00 | 0.00 | 0.00 | 0.00 | 0.00 | 0.52 | 0.00 | 0.56 | 0.23 | 0.00 | 0.00 | 0.00 |
| | Patient20 | 0.00 | 0.19 | 0.00 | 0.00 | 0.00 | 0.00 | 0.00 | 0.00 | 0.00 | 0.00 | 0.00 | 0.00 | 0.00 | 0.23 | 0.00 | 0.20 | 0.45 | 0.40 | 0.28 | 0.15 | 0.00 | 0.00 |
| | Patient21 | 0.00 | 0.00 | 0.00 | 0.00 | 0.00 | 0.00 | 0.15 | 0.00 | 0.20 | 0.00 | 0.00 | 0.00 | 0.00 | 0.14 | 0.00 | 0.00 | 0.13 | 0.12 | 0.42 | 0.00 | 0.00 | 0.00 |
| | Patient22 | 0.00 | 0.00 | 0.00 | 0.00 | 0.25 | 0.00 | 0.09 | 0.00 | 0.29 | 0.11 | 0.00 | 0.00 | 0.00 | 0.30 | 0.00 | 0.00 | 0.30 | 0.52 | 0.33 | 0.00 | 0.00 | 0.00 |
| | Patient23 | 0.64 | 0.36 | 0.00 | 0.00 | 0.19 | 0.00 | 0.00 | 0.00 | 0.00 | 0.00 | 0.00 | 0.00 | 0.00 | 0.21 | 0.01 | 0.00 | 0.08 | 0.42 | 0.19 | 0.00 | 0.00 | 0.00 |
| | Patient24 | 0.00 | 0.20 | 0.00 | 0.00 | 0.00 | 0.00 | 0.00 | 0.00 | 0.00 | 0.00 | 0.00 | 0.00 | 0.11 | 0.09 | 0.00 | 0.00 | 0.06 | 0.82 | 0.53 | 0.00 | 0.00 | 0.00 |
| | Patient25 | 0.00 | 0.00 | 0.00 | 0.00 | 0.00 | 0.00 | 0.00 | 0.00 | 0.38 | 0.00 | 0.00 | 0.00 | 0.00 | 0.26 | 0.00 | 0.00 | 0.00 | 0.18 | 0.09 | 0.00 | 0.00 | 0.00 |
| | Patient26 | 0.40 | 0.00 | 0.00 | 0.00 | 0.00 | 0.00 | 0.09 | 0.00 | 0.00 | 0.00 | 0.00 | 0.00 | 0.25 | 0.04 | 0.00 | 0.00 | 0.07 | 0.38 | 0.13 | 0.46 | 0.00 | 0.03 |
| | Patient27 | 0.00 | 0.00 | 0.00 | 0.43 | 0.00 | 0.00 | 0.00 | 0.00 | 0.00 | 0.00 | 0.00 | 0.00 | 0.00 | 0.09 | 0.00 | 0.05 | 0.10 | 0.22 | 0.00 | 0.00 | 0.00 | 0.00 |
| | Patient28 | 0.00 | 0.00 | 0.33 | 0.00 | 0.00 | 0.00 | 0.00 | 0.00 | 0.00 | 0.00 | 0.00 | 0.00 | 0.00 | 0.00 | 0.00 | 0.19 | 0.00 | 0.00 | 0.00 | 0.00 | 0.00 | 0.00 |
| | Patient29 | 1.99 | 0.28 | 0.00 | 0.54 | 0.25 | 0.02 | 0.00 | 0.00 | 0.10 | 0.00 | 0.00 | 0.00 | 0.00 | 0.10 | 0.00 | 0.00 | 0.00 | 1.06 | 0.33 | 0.00 | 0.00 | 0.00 |
| | Patient30 | 0.00 | 0.00 | 0.00 | 0.80 | 0.00 | 0.24 | 0.00 | 0.00 | 0.15 | 0.00 | 0.00 | 0.00 | 0.00 | 0.15 | 0.04 | 0.00 | 0.06 | 0.66 | 0.08 | 0.00 | 0.00 | 0.69 |

| | | | | | | | | | | | | | | | | | | | | | | | | | | | | | | | | | | | | | | | | | | | | | | | | | | | | | | | | | |
|---|---|---|---|---|---|---|---|---|---|---|---|---|---|---|---|---|---|---|---|---|---|---|---|---|---|---|---|---|---|---|---|---|---|---|---|---|---|---|---|---|---|---|---|---|---|---|---|---|---|---|---|---|---|---|---|---|---|
| PA1 | 0.85 | 0.00 | 0.00 | 0.00 | 0.00 | 0.00 | 0.00 | 0.00 | 0.17 | 0.00 | 0.00 | 0.00 | 0.28 | 0.00 | 0.00 | 0.00 | 0.57 | 0.00 | 0.00 | 0.00 | 0.00 | 0.00 | 0.00 | 0.00 | 0.00 | 0.00 | 0.00 | 0.00 | 0.01 | 0.00 | 0.00 | 0.00 | 0.00 | 0.00 | 0.00 | 0.00 | 0.00 | 0.00 | 0.00 | 0.00 | 0.00 | 0.00 | 0.00 | 0.00 | 0.00 | 0.00 | 0.00 | 0.00 | 0.12 | 0.09 | 0.08 | 0.08 | 0.16 | 0.08 | 0.00 | 0.00 | 0.00 |
| PA2 | 0.86 | 0.00 | 0.01 | 0.00 | 0.00 | 0.00 | 0.00 | 0.00 | 0.09 | 0.00 | 0.00 | 0.00 | 0.00 | 0.00 | 0.00 | 0.00 | 0.14 | 0.00 | 0.00 | 0.00 | 0.00 | 0.00 | 0.00 | 0.00 | 0.00 | 0.00 | 0.00 | 0.00 | 0.00 | 0.00 | 0.00 | 0.00 | 0.00 | 0.00 | 0.00 | 0.08 | 0.25 | 0.00 | 0.00 | 0.00 | 0.00 | 0.00 | 0.00 | 0.00 | 0.00 | 0.00 | 0.00 | 0.00 | 0.00 | 0.18 | 0.00 | 0.39 | 0.06 | 0.25 | 0.00 | 0.00 | 0.00 |
| PA3 | 0.15 | 0.00 | 0.00 | 0.50 | 0.13 | 0.00 | 0.00 | 0.00 | 0.00 | 0.00 | 0.00 | 0.00 | 0.00 | 0.00 | 0.00 | 0.00 | 0.31 | 0.10 | 0.00 | 0.00 | 0.00 | 0.00 | 0.00 | 0.00 | 0.00 | 0.00 | 0.00 | 0.00 | 0.00 | 0.00 | 0.00 | 0.00 | 0.00 | 0.00 | 0.00 | 0.00 | 0.00 | 0.00 | 0.00 | 0.00 | 0.00 | 0.00 | 0.00 | 0.00 | 0.00 | 0.00 | 0.00 | 0.00 | 0.07 | 0.00 | 0.00 | 0.00 | 0.27 | 0.00 | 0.00 | 0.00 | 0.00 |
| PA4 | 0.13 | 0.00 | 0.00 | 0.00 | 0.00 | 0.00 | 0.00 | 0.00 | 0.18 | 0.00 | 0.00 | 0.00 | 0.00 | 0.00 | 0.00 | 0.69 | 0.26 | 0.00 | 0.00 | 0.00 | 0.00 | 0.00 | 0.00 | 0.00 | 0.00 | 0.00 | 0.00 | 0.00 | 0.00 | 0.00 | 0.00 | 0.00 | 0.00 | 0.00 | 0.00 | 0.00 | 0.00 | 0.00 | 0.00 | 0.00 | 0.00 | 0.00 | 0.00 | 0.00 | 0.00 | 0.00 | 0.00 | 0.00 | 0.08 | 0.00 | 0.00 | 0.00 | 0.10 | 0.00 | 0.00 | 0.00 | 0.49 |
| PA5 | 0.31 | 0.00 | 0.01 | 0.32 | 0.25 | 0.00 | 0.03 | 0.00 | 0.10 | 0.00 | 0.00 | 0.00 | 0.00 | 0.00 | 0.00 | 0.00 | 0.36 | 0.00 | 0.00 | 0.00 | 0.00 | 0.00 | 0.00 | 0.00 | 0.00 | 0.00 | 0.00 | 0.00 | 0.00 | 0.00 | 0.00 | 0.00 | 0.00 | 0.00 | 0.00 | 0.00 | 0.00 | 0.00 | 0.00 | 0.00 | 0.00 | 0.00 | 0.00 | 0.00 | 0.00 | 0.00 | 0.00 | 0.00 | 0.00 | 0.00 | 0.00 | 0.09 | 0.23 | 0.19 | 0.00 | 0.00 | 0.00 |
| PA6 | 0.00 | 0.00 | 0.65 | 0.10 | 0.00 | 0.00 | 0.00 | 0.00 | 0.15 | 0.00 | 0.00 | 0.00 | 0.00 | 0.00 | 0.00 | 0.00 | 0.23 | 0.00 | 0.00 | 0.00 | 0.00 | 0.00 | 0.00 | 0.00 | 0.00 | 0.00 | 0.00 | 0.00 | 0.00 | 0.00 | 0.00 | 0.00 | 0.00 | 0.00 | 0.27 | 0.00 | 0.00 | 0.00 | 0.00 | 0.00 | 0.00 | 0.00 | 0.00 | 0.00 | 0.00 | 0.00 | 0.00 | 0.00 | 0.00 | 0.00 | 0.00 | 0.07 | 0.21 | 0.07 | 0.00 | 0.00 | 0.00 |
| PA7 | 1.22 | 0.00 | 0.00 | 0.00 | 0.10 | 0.00 | 1.21 | 0.00 | 0.92 | 0.00 | 0.35 | 0.00 | 0.00 | 0.82 | 0.00 | 0.00 | 0.88 | 0.00 | 0.00 | 0.00 | 0.00 | 0.00 | 0.00 | 0.00 | 0.00 | 0.00 | 0.00 | 0.00 | 0.00 | 0.00 | 0.00 | 0.00 | 0.21 | 0.00 | 0.09 | 0.00 | 0.00 | 0.00 | 0.00 | 0.00 | 0.00 | 0.00 | 0.00 | 0.00 | 0.00 | 0.00 | 0.00 | 0.00 | 0.17 | 0.00 | 0.00 | 0.10 | 0.59 | 0.09 | 0.00 | 0.00 | 0.00 |
| PA8 | 0.88 | 0.00 | 0.00 | 0.00 | 0.00 | 0.00 | 0.00 | 0.00 | 0.14 | 0.00 | 0.00 | 0.00 | 0.32 | 0.00 | 0.00 | 0.00 | 0.84 | 0.00 | 0.00 | 0.00 | 0.00 | 0.28 | 0.00 | 0.00 | 0.20 | 0.00 | 0.00 | 0.00 | 0.00 | 0.00 | 0.00 | 0.00 | 0.00 | 0.24 | 0.06 | 0.00 | 0.11 | 0.00 | 0.00 | 0.00 | 0.00 | 0.00 | 0.00 | 0.00 | 0.00 | 0.00 | 0.00 | 0.00 | 0.10 | 0.29 | 0.00 | 0.00 | 0.51 | 0.16 | 0.00 | 0.00 | 0.00 |
| PA9 | 1.28 | 0.00 | 0.00 | 0.00 | 0.00 | 0.00 | 0.00 | 0.00 | 0.31 | 0.00 | 0.00 | 0.00 | 0.00 | 0.00 | 0.00 | 0.00 | 1.43 | 0.00 | 0.00 | 0.00 | 0.00 | 0.00 | 0.00 | 0.00 | 0.00 | 0.00 | 0.00 | 0.00 | 0.00 | 0.00 | 0.00 | 0.00 | 0.00 | 0.00 | 0.00 | 1.54 | 0.00 | 0.00 | 0.00 | 0.00 | 0.00 | 0.00 | 0.00 | 0.00 | 0.00 | 0.00 | 0.00 | 0.00 | 0.08 | 0.12 | 0.00 | 0.12 | 0.44 | 0.43 | 0.00 | 0.00 | 0.34 |
| PA10 | 1.01 | 0.00 | 0.00 | 0.00 | 0.03 | 0.00 | 0.00 | 0.00 | 0.82 | 0.00 | 0.00 | 0.00 | 0.00 | 0.00 | 0.00 | 0.00 | 1.24 | 0.00 | 0.00 | 0.00 | 0.00 | 0.00 | 0.00 | 0.00 | 0.00 | 0.00 | 0.00 | 0.00 | 0.00 | 0.00 | 0.00 | 0.00 | 0.00 | 0.15 | 0.40 | 0.00 | 0.00 | 0.00 | 0.32 | 0.00 | 0.00 | 0.00 | 0.00 | 0.00 | 0.00 | 0.00 | 0.00 | 0.00 | 0.08 | 0.00 | 0.00 | 0.08 | 0.42 | 0.09 | 0.00 | 0.00 | 0.00 |

*This transcript is not in the current gene set.

**b.**

| | Protein coding | | | | | | | | | | | | | | | |
|---|---|---|---|---|---|---|---|---|---|---|---|---|---|---|---|---|
| | ATXN3-251 (ref.) | ATXN3-201 | ATXN3-203 | ATXN3-204 | ATXN3-206 | ATXN3-207 | ATXN3-209 | ATXN3-213 | ATXN3-214 | ATXN3-215 | ATXN3-219 | ATXN3-227 | ATXN3-228 | ATXN3-231 | ATXN3-235 | ATXN3-246 |
| Cerebellum | 10 | 11 | 3 | 1 | 7 | 8 | 1 | 2 | 12 | 0 | 11 | 7 | 0 | 2 | 4 | 1 |
| | 83% | 92% | 25% | 8% | 58% | 67% | 8% | 17% | 100% | 0% | 92% | 58% | 0% | 17% | 33% | 8% |
| Controls, n=6 | 5 | 6 | 2 | 1 | 5 | 4 | 1 | 1 | 6 | 0 | 6 | 4 | 0 | 2 | 1 | 1 |
| | 83% | 100% | 33% | 17% | 83% | 67% | 17% | 17% | 100% | 0% | 100% | 67% | 0% | 33% | 17% | 17% |
| Patients, n=6 | 5 | 5 | 1 | 0 | 2 | 4 | 0 | 1 | 6 | 0 | 5 | 3 | 0 | 0 | 3 | 0 |
| | 83% | 83% | 17% | 0% | 33% | 67% | 0% | 17% | 100% | 0% | 83% | 50% | 0% | 0% | 50% | 0% |
| Blood | 58 | 6 | 15 | 6 | 11 | 6 | 13 | 5 | 45 | 1 | 10 | 3 | 12 | 5 | 5 | 7 |
| | 97% | 10% | 25% | 10% | 18% | 10% | 22% | 8% | 75% | 2% | 17% | 5% | 20% | 8% | 8% | 12% |
| Controls, n=20 | 20 | 2 | 6 | 0 | 2 | 5 | 7 | 1 | 16 | 1 | 3 | 1 | 3 | 1 | 3 | 2 |
| | 100% | 10% | 30% | 0% | 10% | 25% | 35% | 5% | 80% | 5% | 15% | 5% | 15% | 5% | 15% | 10% |
| Patients, n=30 | 29 | 4 | 6 | 3 | 5 | 1 | 4 | 4 | 20 | 0 | 6 | 2 | 7 | 3 | 2 | 4 |
| | 97% | 13% | 20% | 10% | 17% | 3% | 13% | 13% | 67% | 0% | 20% | 7% | 23% | 10% | 7% | 13% |
| PA, n=10 | 9 | 0 | 3 | 3 | 4 | 0 | 2 | 0 | 9 | 0 | 1 | 0 | 2 | 1 | 0 | 1 |
| | 90% | 0% | 30% | 30% | 40% | 0% | 20% | 0% | 90% | 0% | 10% | 0% | 20% | 10% | 0% | 10% |
| *ATXN3* mutation carriers, n=40 | 38 | 4 | 9 | 6 | 9 | 1 | 6 | 4 | 29 | 0 | 7 | 2 | 9 | 4 | 2 | 5 |
| | 95% | 10% | 23% | 15% | 23% | 3% | 15% | 10% | 73% | 0% | 18% | 5% | 23% | 10% | 5% | 13% |

| | Nonsense mediated decay | | | | | | | | | | | | | | | | | | |
|---|---|---|---|---|---|---|---|---|---|---|---|---|---|---|---|---|---|---|---|
| | ATXN3-202 | ATXN3-212 | ATXN3-218 | ATXN3-221 | ATXN3-225 | ATXN3-229 | ATXN3-230 | ATXN3-232 | ATXN3-234 | ATXN3-236 | ATXN3-237 | ATXN3-238 | ATXN3-240 | ATXN3-242 | ATXN3-243 | ATXN3-244 | ATXN3-245 | ATXN3-253 | ATXN3-254 |
| Cerebellum | 4 | 0 | 0 | 0 | 6 | 1 | 4 | 3 | 11 | 0 | 4 | 3 | 5 | 1 | 9 | 4 | 3 | 8 | 8 |
| | 33% | 0% | 0% | 0% | 50% | 8% | 33% | 25% | 92% | 0% | 33% | 25% | 42% | 8% | 75% | 33% | 25% | 67% | 67% |
| Controls, n=6 | 3 | 0 | 0 | 0 | 3 | 1 | 3 | 1 | 6 | 0 | 3 | 1 | 3 | 0 | 4 | 3 | 1 | 4 | 4 |
| | 50% | 0% | 0% | 0% | 50% | 17% | 50% | 17% | 100% | 0% | 50% | 17% | 50% | 0% | 67% | 50% | 17% | 67% | 67% |
| Patients, n=6 | 1 | 0 | 0 | 0 | 3 | 0 | 1 | 2 | 5 | 0 | 1 | 2 | 2 | 1 | 5 | 1 | 2 | 4 | 4 |
| | 17% | 0% | 0% | 0% | 50% | 0% | 17% | 33% | 83% | 0% | 17% | 33% | 33% | 17% | 83% | 17% | 33% | 67% | 67% |
| Blood | 59 | 3 | 1 | 2 | 1 | 14 | 1 | 2 | 8 | 1 | 7 | 3 | 1 | 0 | 6 | 0 | 1 | 10 | 31 |
| | 98% | 5% | 2% | 3% | 2% | 23% | 2% | 3% | 13% | 2% | 12% | 5% | 2% | 0% | 10% | 0% | 2% | 17% | 52% |
| Controls, n=20 | 20 | 1 | 1 | 1 | 0 | 7 | 1 | 0 | 5 | 0 | 4 | 1 | 0 | 0 | 2 | 0 | 0 | 4 | 8 |
| | 100% | 5% | 5% | 5% | 0% | 35% | 5% | 0% | 25% | 0% | 20% | 5% | 0% | 0% | 10% | 0% | 0% | 20% | 40% |
| Patients, n=30 | 29 | 1 | 0 | 1 | 1 | 6 | 0 | 2 | 2 | 1 | 3 | 2 | 0 | 0 | 4 | 0 | 0 | 4 | 19 |
| | 97% | 3% | 0% | 3% | 3% | 20% | 0% | 7% | 7% | 3% | 10% | 7% | 0% | 0% | 13% | 0% | 0% | 13% | 63% |
| PA, n=10 | 10 | 1 | 0 | 0 | 0 | 1 | 0 | 0 | 1 | 0 | 0 | 0 | 1 | 0 | 0 | 0 | 1 | 2 | 4 |
| | 100% | 10% | 0% | 0% | 0% | 10% | 0% | 0% | 10% | 0% | 0% | 0% | 10% | 0% | 0% | 0% | 10% | 20% | 40% |
| *ATXN3* mutation carriers, n=40 | 39 | 2 | 0 | 1 | 1 | 7 | 0 | 2 | 3 | 1 | 3 | 2 | 1 | 0 | 4 | 0 | 1 | 6 | 23 |
| | 98% | 5% | 0% | 3% | 3% | 18% | 0% | 5% | 8% | 3% | 8% | 5% | 3% | 0% | 10% | 0% | 3% | 15% | 58% |

| | Protein coding CDS not defined | | | | | | | | | | | | | | | |
|---|---|---|---|---|---|---|---|---|---|---|---|---|---|---|---|---|
| | ATXN3-208 | ATXN3-211 | ATXN3-216 | ATXN3-217 | ATXN3-220 | ATXN3-222 | ATXN3-223 | ATXN3-224 | ATXN3-226 | ATXN3-233 | ATXN3-239 | ATXN3-241 | ATXN3-247 | ATXN3-249 | ATXN3-250 | ATXN3-252 |
| Cerebellum | 12 | 2 | 1 | 10 | 0 | 1 | 2 | 2 | 8 | 0 | 1 | 1 | 1 | 1 | 2 | 1 |
| | 100% | 17% | 8% | 83% | 0% | 8% | 17% | 17% | 67% | 0% | 8% | 8% | 8% | 8% | 17% | 8% |
| Controls, n=6 | 6 | 2 | 1 | 5 | 0 | 0 | 0 | 1 | 6 | 0 | 1 | 1 | 0 | 0 | 1 | 0 |
| | 100% | 33% | 17% | 83% | 0% | 0% | 0% | 17% | 100% | 0% | 17% | 17% | 0% | 0% | 17% | 0% |
| Patients, n=6 | 6 | 0 | 0 | 5 | 0 | 1 | 2 | 1 | 2 | 0 | 0 | 0 | 1 | 1 | 1 | 1 |
| | 100% | 0% | 0% | 83% | 0% | 17% | 33% | 17% | 33% | 0% | 0% | 0% | 17% | 17% | 17% | 17% |
| Blood | 14 | 16 | 5 | 7 | 8 | 4 | 5 | 7 | 13 | 8 | 1 | 2 | 10 | 41 | 15 | 8 |
| | 23% | 27% | 8% | 12% | 13% | 7% | 8% | 12% | 22% | 13% | 2% | 3% | 17% | 68% | 25% | 13% |
| Controls, n=20 | 3 | 5 | 1 | 0 | 4 | 0 | 0 | 4 | 6 | 3 | 1 | 1 | 3 | 12 | 7 | 1 |
| | 15% | 25% | 5% | 0% | 20% | 0% | 0% | 20% | 30% | 15% | 5% | 5% | 15% | 60% | 35% | 5% |
| Patients, n=30 | 9 | 9 | 4 | 6 | 4 | 4 | 5 | 3 | 7 | 5 | 0 | 1 | 7 | 22 | 4 | 6 |
| | 30% | 30% | 13% | 20% | 13% | 13% | 17% | 10% | 23% | 17% | 0% | 3% | 23% | 73% | 13% | 20% |
| PA, n=10 | 2 | 2 | 0 | 1 | 0 | 0 | 0 | 0 | 0 | 0 | 0 | 0 | 0 | 7 | 4 | 1 |
| | 20% | 20% | 0% | 10% | 0% | 0% | 0% | 0% | 0% | 0% | 0% | 0% | 0% | 70% | 40% | 10% |
| *ATXN3* mutation carriers, n=40 | 11 | 11 | 4 | 7 | 4 | 4 | 5 | 3 | 7 | 5 | 0 | 1 | 7 | 29 | 8 | 7 |
| | 28% | 28% | 10% | 18% | 10% | 10% | 13% | 8% | 18% | 13% | 0% | 3% | 18% | 73% | 20% | 18% |

| | Retained intron | | |
|---|---|---|---|
| | ATXN3-205 | ATXN3-210 | ATXN3-248 |
| Cerebellum | 12 | 9 | 2 |
| | 100% | 75% | 17% |
| Controls, n=6 | 6 | 4 | 1 |
| | 100% | 67% | 17% |
| Patients, n=6 | 6 | 5 | 1 |
| | 100% | 83% | 17% |
| Blood | 35 | 52 | 42 |
| | 58% | 87% | 70% |
| Controls, n=20 | 12 | 17 | 16 |
| | 60% | 85% | 80% |
| Patients, n=30 | 16 | 25 | 18 |
| | 53% | 83% | 60% |
| PA, n=10 | 7 | 10 | 8 |
| | 70% | 100% | 80% |
| *ATXN3* mutation carriers, n=40 | 23 | 35 | 26 |
| | 58% | 88% | 65% |

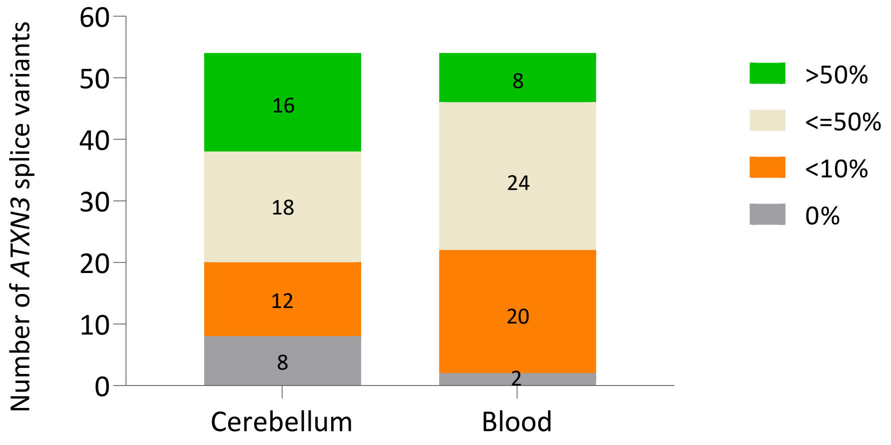


**Supplementary figure 1.** Number of transcripts by frequency in the 72 samples (12 samples from cerebellum and the 60 samples from blood).

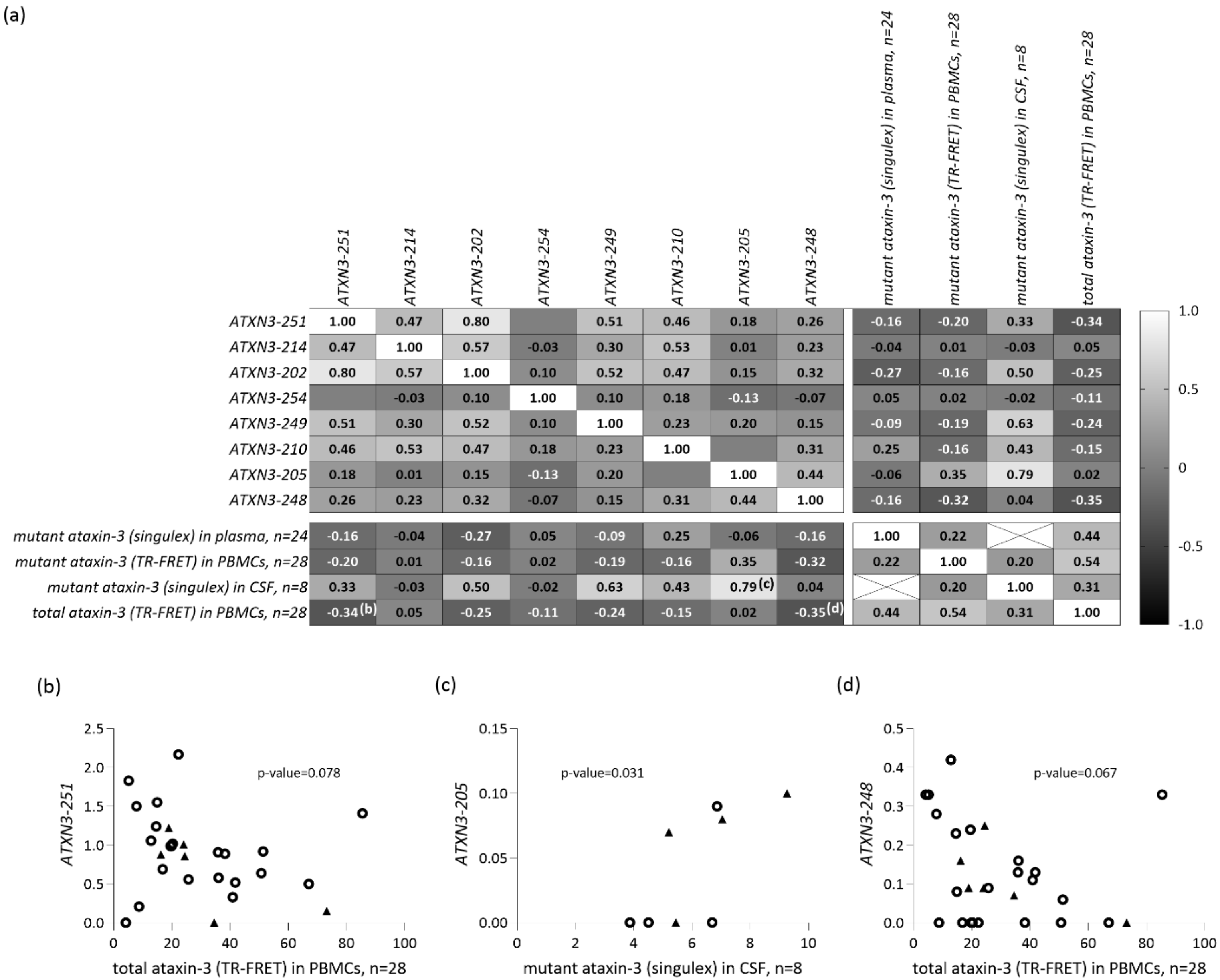

(a)
ATXN3-251
ATXN3-214
ATXN3-202
ATXN3-254
ATXN3-249
ATXN3-210
ATXN3-205
ATXN3-248
mutant ataxin-3 (singulex) in plasma, n=24
mutant ataxin-3 (TR-FRET) in PBMCs, n=28
mutant ataxin-3 (singulex) in CSF, n=8
total ataxin-3 (TR-FRET) in PBMCs, n=28
ATXN3-251 1.00 0.47 0.80 0.51 0.46 0.18 0.26 -0.16 -0.20 0.33 -0.34
ATXN3-214 0.47 1.00 0.57 -0.03 0.30 0.53 0.01 0.23 -0.04 0.01 -0.03 0.05
ATXN3-202 0.80 0.57 1.00 0.10 0.52 0.47 0.15 0.32 -0.27 -0.16 0.50 -0.25
ATXN3-254 -0.03 0.10 1.00 0.10 0.18 -0.13 -0.07 0.05 0.02 -0.02 -0.11
ATXN3-249 0.51 0.30 0.52 0.10 1.00 0.23 0.20 0.15 -0.09 -0.19 0.63 -0.24
ATXN3-210 0.46 0.53 0.47 0.18 0.23 1.00 0.31 0.25 -0.16 0.43 -0.15
ATXN3-205 0.18 0.01 0.15 -0.13 0.20 1.00 0.44 -0.06 0.35 0.79 0.02
ATXN3-248 0.26 0.23 0.32 -0.07 0.15 0.31 0.44 1.00 -0.16 -0.32 0.04 -0.35
mutant ataxin-3 (singulex) in plasma, n=24 -0.16 -0.04 -0.27 0.05 -0.09 0.25 -0.06 -0.16 1.00 0.22 0.44
mutant ataxin-3 (TR-FRET) in PBMCs, n=28 -0.20 0.01 -0.16 0.02 -0.19 -0.16 0.35 -0.32 0.22 1.00 0.20 0.54
mutant ataxin-3 (singulex) in CSF, n=8 0.33 -0.03 0.50 -0.02 0.63 0.43 0.79 (c) 0.04 0.20 1.00 0.31
total ataxin-3 (TR-FRET) in PBMCs, n=28 -0.34 (b) 0.05 -0.25 -0.11 -0.24 -0.15 0.02 -0.35 (d) 0.44 0.54 0.31 1.00
1.0
0.5
0
-0.5
-1.0
(b)
ATXN3-251
p-value=0.078
total ataxin-3 (TR-FRET) in PBMCs, n=28
(c)
ATXN3-205
p-value=0.031
mutant ataxin-3 (singulex) in CSF, n=8
(d)
ATXN3-248
p-value=0.067
total ataxin-3 (TR-FRET) in PBMCs, n=28

**Supplementary Figure 2.** Spearman rank correlation matrix (a) of the levels of eight *ATXN3* splice variants frequently expressed in blood with ataxin-3 protein (mutant and total) levels measured in plasma, peripheral mononuclear blood cells (PBMCs) and cerebrospinal fluid (CSF), using two different methodologies (Singulex and TR-FRET)[a,b]. For transcripts whose p-value were lower than 0.05 or borderline after a Spearman rank correlation test, the individual values were shown in detail (b), (c) and (d). In the graphs, SCA3/MJD patients are represented as **O** and pre-ataxic carriers as ▲.

a. Gonsior K, Kaucher GA, Pelz P, Schumann D, Gansel M, Kuhs S, et al. PolyQ-expanded ataxin-3 protein levels in peripheral blood mononuclear cells correlate with clinical parameters in SCA3: a pilot study. J Neurol. 2021 Apr 26;268(4):1304–15.
b. Hübener-Schmid J, Kuhlbrodt K, Peladan J, Faber J, Santana MM, Hengel H, et al. Polyglutamine-Expanded Ataxin-3: A Target Engagement Marker for Spinocerebellar Ataxia Type 3 in Peripheral Blood. Mov Disord. 2021 Nov 16;36(11):2675–81.

**Supplementary table 3.** Additional members of the European Spinocerebellar Ataxia type 3/Machado-Joseph Initiative (ESMI) Study Group.

| Site | Name | Surname | Degree | Institution | Department | City | Country | E-mail |
|---|---|---|---|---|---|---|---|---|
| Aachen | Janna | Krahe | | RWTH Aachen University | Neurology | Aachen | Germany | |
| | Kathrin | Reetz | MD | RWTH Aachen University | Neurology | Aachen | Germany | |
| Azores | Pedro | Lopes | MD | Hospital do Divino Espírito Santo | Neurology | Ponta Delgada | Portugal | |
| | José | González | MD | Faculdade de Ciências e Tecnologia, Universidade dos Açores | | Ponta Delgada | Portugal | |
| | Carlos | Gonzalez | BSc | Faculdade de Ciências e Tecnologia, Universidade dos Açores | | Ponta Delgada | Portugal | |
| | João | Lemos | BSc | Hospital do Santo Espírito da Ilha Terceira | Departamento de Psiquiatria e Saúde Mental | Angra do Heroísmo | Portugal | |
| Bonn | Ilaria | Giordano | MD | 1. German Center for Neurodegenerative Diseases (DZNE), Bonn<br>2. University Hospital of Bonn | 1. N/A<br>2. Department of Neurology | Bonn | Germany | |
| | Marcus | Grobe-Einsler | MD | 1. German Center for Neurodegenerative Diseases (DZNE), Bonn<br>2. University Hospital of Bonn | 1. N/A<br>2. Department of Neurology | Bonn | Germany | |
| | Demet | Önder | MD | 1. German Center for Neurodegenerative Diseases (DZNE), Bonn<br>2. University Hospital of Bonn | 1. N/A<br>2. Department of Neurology | Bonn | Germany | |
| Coimbra | Patrick | Silva | MSc | CNC - Center for Neuroscience and Cell Biology, University of Coimbra, Coimbra, Portugal;<br>CIBB- Center for Innovation in Biomedicine and Biotechnology, University of Coimbra, Coimbra, Portugal;<br>IIIUC - Institute for Interdisciplinary Research, University of Coimbra, Coimbra, Portugal; | | Coimbra | Portugal | |
| | Cristina | Januário | MD, PhD | Faculty of Medicine, University of Coimbra, Coimbra, Portugal | | Coimbra | Portugal | |
| | Joana | Ribeiro | MD | Child Development Centre, Pediatric Hospital, Coimbra University Hospital Centre, University of Coimbra, Coimbra, Portugal | | Coimbra | Portugal | |

| | | | | | | | | |
|---|---|---|---|---|---|---|---|---|
| | Inês | Cunha | MD | Neurology unit, Coimbra University Hospital Centre, University of Coimbra, Coimbra, Portugal. | | Coimbra | Portugal | |
| | João | Lemos | MD, PhD | Coimbra University Hospital Centre, University of Coimbra, Coimbra, Portugal. | | Coimbra | Portugal | |
| | Maria M | Pinto | MSc | Center for Neuroscience and Cell Biology (CNC), University of Coimbra, 3004-504 Coimbra, Portugal;<br>CIBB- Center for Innovation in Biomedicine and Biotechnology, University of Coimbra, Coimbra, Portugal; | | Coimbra | Portugal | |
| Essen | Dagmar | Timmann | MD | Essen University Hospital | Department of Neurology | Essen | Germany | |
| | Katharina M. | Steiner | MD | Essen University Hospital | Department of Neurology | Essen | Germany | |
| | Andreas | Thieme | MD | Essen University Hospital | Department of Neurology | Essen | Germany | |
| | Thomas M. | Ernst | MSc | Essen University Hospital | Department of Neurology | Essen | Germany | |
| Heidelberg | Heike | Jacobi | MD | Heidelberg University Hospital | Department of Neurology | Heidelberg | Germany |-heidelberg.d |
| London | Nita | Solanky | PhD | University College London/National Hospital for Neurology and Neurosurgery, University College London Hospitals NHS Foundation Trust | Department of Clinical and Movement Neurosciences | London | United Kingdom | |
| | Cristina | Gonzalez-Robles | MD | University College London/National Hospital for Neurology and Neurosurgery, University College London Hospitals NHS Foundation Trust | Department of Clinical and Movement Neurosciences | London | United Kingdom | |
| Nijmegen | Judith | Van Gaalen | MD | Donders Institute for Brain, Cognition, and Behavior, Radboud university medical center | Expert centre for Parkinson & Movement Disorders, Department of Neurology | Nijmegen | The Netherlands | |
| Santander | Ana Lara | Pelayo-Negro | MD, PhD | University Hospital Marqués de Valdecilla-IDIVAL | Neurology | Santander | Spain | |
| | Leire | Manrique | MD | University Hospital Marqués de Valdecilla-IDIVAL | Neurology | Santander | Spain | |

| | | | | | | | | |
|---|---|---|---|---|---|---|---|---|
| Tübingen | Holger | Hengel | MD | University of Tübingen | Department of Neurology and Hertie Institute for Clinical Brain Research | Tübingen | Germany | |
| | Matthis | Synofzik | MD | University of Tübingen | Department of Neurology and Hertie Institute for Clinical Brain Research | Tübingen | Germany | |
| | Winfried | Ilg | PhD | University of Tübingen | Hertie Institute for Clinical Brain Research | Tübingen | Germany | |
